# mTORC1 signaling modulate microtubule tyrosination/detyrosination status to regulate lysosome dynamics

**DOI:** 10.1101/2025.09.21.677577

**Authors:** Jeganath Ammavasai, Nitin Mohan

## Abstract

Lysosomes integrate nutrient sensing and metabolic signaling by dynamically repositioning within the cytoplasm. While motors and lysosomal surface adaptors driving this movement are well studied, how the microtubule cytoskeleton specifies lysosomal transport and its coupling to mechanistic target of rapamycin complex 1 (mTORC1) signaling remains unclear. Here, we show that tubulin post-translational modifications (PTMs) define the tracks that guide nutrient-responsive lysosome positioning. Using super-resolution imaging and biochemical assays, we demonstrate that lysosomes carrying active mTORC1 preferentially traffic along tyrosinated microtubules to reach the cell periphery, whereas detyrosinated microtubules restrict their motility. Nutrient stimulation enhances this outward transport, while concurrently suppressing microtubule detyrosination through mTORC1-S6K1 signaling, likely via inhibition of vasohibin activity. This ensures selective expansion of tyrosinated microtubule tracks to sustain mTORC1 signaling. Strikingly, in cancer cells where aberrant mTORC1 activity correlates with altered microtubule tyrosination, restoring microtubule detyrosination rescues mTORC1 hyperactivation and suppresses cell migration and epithelial-to-mesenchymal transition. Thus, demonstrating that tubulin post-translational modifications (PTMs) define the tracks that guide the positioning of nutrient-responsive lysosomes. These findings uncover a crosstalk between mTORC1 and tubulin tyrosination/detyrosination status that couples nutrient availability to lysosomal dynamics and identify microtubule tyrosination as a regulatory node with therapeutic potential in cancer.

## Introduction

Lysosomes are recognized as the cell’s primary degradation centers, receiving cargo from endocytic, phagocytic, and autophagic pathways. Lysosomal enzymes maintain quality control through the proper degradation and recycling of materials received from both within and outside the cell. Recent research, however, indicates that lysosomes are more than just recycling sites; they are essential hubs for sensing cellular nutrients and regulating metabolic signals (Sancak et al., 2010). The lysosomal membrane features transmembrane proteins and ion channels that detect nutrient levels and control the storage or exchange of metabolites between the lysosomal lumen and the cytosol. In nutrient-rich conditions, the mechanistic target of rapamycin complex 1 (mTORC1), a kinase complex, which senses cellular nutrients, moves to the lysosomal surface, becomes active (Menon et al., 2014; Sancak et al., 2010), and promotes cell growth and proliferation. In contrast, during nutrient scarcity, TSC1/2 proteins localize to lysosomes and suppress mTORC1 activity (Demetriades et al., 2016; Menon et al., 2014). When mTORC1 is inactive, it disassociates from the lysosomal surface (Sancak et al., 2010), enabling lysosomes to engage in autophagy. Lysosomes thus play a central role in directing the cell towards anabolic or catabolic states based on nutrient availability.

Emerging evidence shows that lysosomes adapt their physical and chemical properties in response to fluctuating nutrient levels, likely to accommodate conflicting molecular complexes and perform opposing functions. During starvation, lysosomes increase in size (500-1500 nm in diameter) and decrease in number within the cell (Yu et al., 2010)(Bandyopadhyay et al., 2014). Their luminal pH becomes more acidic, and they tend to cluster near the nucleus (Korolchuk et al., 2011), promoting their fusion with autophagosomes. Conversely, lysosomes are more dispersed throughout the cytoplasm, with hundreds present per cell, measuring 100-500 nm in diameter, during fed states (Mellman, 1989). Additionally, when cells transition from starved to fed conditions, a lipid switch occurs at the lysosomal membrane from phosphoinositol-4-phosphate (PI(4)P) to phosphoinositol-3-phosphate (PI(3)P) to activate mTORC1 on its surface (Ebner et al., 2023). Furthermore, several studies show a significant shift in lysosomal positioning from the perinuclear region to the cell periphery in response to nutrient stimulation, and inhibiting this lysosomal reorganization suppresses mTORC1 function (Ebner et al., 2023; Jerabkova-Roda et al., 2025; B. Wang et al., 2021).

Such rapid repositioning of lysosomes, critical for nutrient signaling, involves long-distance bidirectional transport steered by motor proteins and microtubules. Kinesins move lysosomes outward toward the cell periphery (Nakata & Hirokawa, 1995; Rosa-Ferreira & Munro, 2011) while dynein directs retrograde movement toward the perinuclear area (Jordens et al., 2001). Changes in lysosomal membrane lipids could trigger the recruitment of specific motors (Rocha et al., 2009), thereby guiding lysosomes to specific cellular locations for distinct functions. The small GTPase Arl8b interacts with SKIP to recruit kinesin motors for peripheral distribution; its increased expression is associated with elevated mTORC1 activity (Guardia et al., 2016; Korolchuk et al., 2011; Rosa-Ferreira & Munro, 2011). Specifically, kinesin-3 family members Kif1A, Kif16B (Blatner et al., 2007; Ebner et al., 2023; Guardia et al., 2016), and kinesin-13 member Kif2A (Korolchuk et al., 2011) facilitate lysosome outward movement, correlating with high mTORC1 activity. Conversely, the transmembrane protein TMEM55B interacts with JIP4, recruiting dynein (Willett et al., 2017) for retrograde transport and promoting fusion with autophagosomes during starvation. Additionally, small GTPase Rab7 interacts with RILP to efficiently recruit dynein, directing lysosomes toward the microtubule minus end (Jordens et al., 2001). Thus, we have a substantial understanding of the coordinated mechanisms on the lysosome surface that recruit specific motors, ensuring their dynamic positioning is attuned to the cell’s metabolic state. However, the mechanisms by which these proteins at the lysosome surface couple them with microtubules and how microtubules control lysosomal transport, positioning and function in relation to nutrient sensing and mTORC1 signalling remain elusive. Tubulin post-translational modifications create different microtubule subpopulations, such as acetylated (L’Hernaul & Rosenbaum, 2002), tyrosinated (Hallak et al., 1977; Raybin & Flavin, 1977), detyrosinated (Aillaud et al., 2017; Kreitzer et al., 1999; Nieuwenhuis et al., 2017), polyglutamylated (Alexander et al., 1991; Eddé et al., 1990), and polyglycylated (Redeker et al., 1994) microtubules, which selectively regulate motor proteins. These modifications control organelle transport, establish membrane contacts between organelles, and are reported to regulate lysosome function (Mohan et al., 2019; Zheng et al., 2021). For example, acetylated microtubules enable autophagosome-lysosome fusion (Xie et al., 2010), while tyrosinated microtubules support LRRK-mediated lysosomal tubulation (Bonet-Ponce et al., 2024) involved in membrane repair. Detyrosinated microtubules have also been shown to immobilize lysosomes to support autophagy (Mohan et al., 2019). These PTMs are linked to microtubule stability. Studies show that microtubule stability and PTM states fluctuate depending on nutrient availability (Knudsen et al., 2023; Parker et al., 2019). For instance, starving HeLa cells pretreated with microtubule stabilizers leads to tubulin hyperacetylation (Geeraert et al., 2010). Furthermore, elevated microtubule acetylation is observed in various cancers (Wattanathamsan et al., 2021; Yoshimoto et al., 2022), correlating with abnormal mTORC1 signalling and uncontrolled cell growth. These findings suggest a connection between tubulin-PTM regulation of lysosomal function and mTORC1 signalling. We propose that microtubule tracks marked by specific tubulin-PTMs facilitate rapid lysosome repositioning in response to nutrients and are essential for mTORC1 signalling.

In this study, by combining super-resolution microscopy and biochemical assays, we demonstrate that, in epithelial cells, tyrosinated microtubules facilitate the outward movement and dispersal of lysosomes containing mTORC1, whereas detyrosinated microtubules limit this transport. Additionally, we find that mTORC1 signalling inhibits microtubule detyrosination via its substrate S6K1, which likely suppresses the detyrosination enzyme vasohibin. Moreover, in certain cancer types where higher tubulin tyrosination levels are linked to abnormal mTORC1 activity, manipulating tyrosination or detyrosination genetically can restore proper mTORC1 activity and control cell migration and epithelial-to-mesenchymal transition, and processes associated with cancer progression.

## Results

### mTORC1 recruitment promotes lysosomal motility

Nutrient signalling localizes mTORC1 on lysosomes. Concurrently, reports also show that nutrient stimulation redistributes lysosomes towards the cell periphery. However, whether mTORC1 recruitment actively drives this lysosomal repositioning remains unclear. To address the impact of mTORC1 on lysosomal motility, we starved BS-C-1 cells for 2 hours, then restimulated them with amino acids and serum, and tracked mTORC1 localization on lysosomes and lysosomal motility at specific time intervals over a 1-hour period (Figure 1A-F). Manual scoring of ∼1000 lysosomes per time point, across four cells, revealed that mTORC1 association with lysosomes peaked at 20 minutes of restimulation (56.4% ± 8.63), nearly double that observed at 10 minutes and 1.5 times higher than at 30 minutes of restimulation (Figure 1A, B). Strikingly, lysosome motility followed a similar trend: the fraction of motile lysosomes rose steadily from 5 minutes, peaked at 25 minutes (53.05% ± 7.06), and declined thereafter (Figure 1C, D). Directional analysis (see Methods and Figure S1A) further revealed a 1.6-fold increase in the percentage of lysosomes moving anterogradely at the peak 25 minutes compared to 5 minutes of restimulation (Figure 1E). These observations demonstrate that nutrient-induced mTORC1 recruitment is followed by enhanced lysosomal motility (Figure 1F), particularly toward the cell periphery.

**Figure 1.**
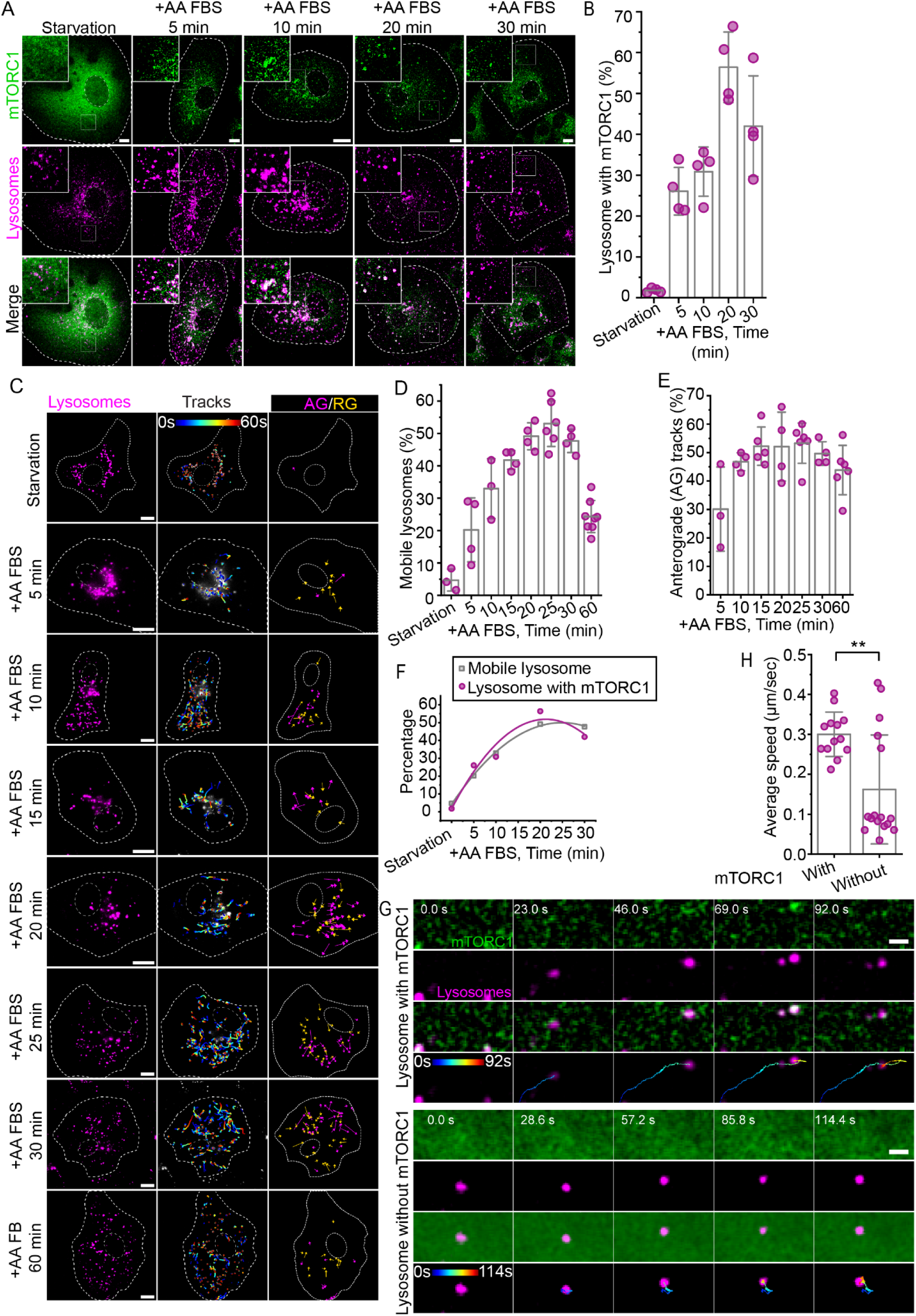
Nutrient stimulation recruits mTORC1 to lysosomes and promotes their motility. **(A)** Two-colour confocal microscopy of endogenous mTOR (green) and LAMP2 (lysosomes, magenta) in BS-C-1 cells starved for 2 hours or starved and restimulated with amino acids and FBS for indicated time points. Insets highlight mTOR puncta colocalized with lysosomes. **(B)** Quantification of lysosomes positive for mTOR signal. Starved cells show minimal mTOR-lysosome colocalization, which increases upon nutrient restimulation, peaking at 20 minutes, followed by a decline at 30 minutes. **(C)** Live-cell TIRF imaging and tracking of lysosomes (Lysotracker) in BS-C-1 cells under starvation and restimulation conditions. Trajectories of individual lysosomes are shown with starting points in blue and endpoints in red. Tracks are categorized into anterograde (AG, magenta) and retrograde (RG, yellow) movement. **(D–E)** Quantification of mobile lysosomes (displacement >1 µm) shows a progressive increase in the mobile population (**D**) and in anterograde movement (**E**) with nutrient stimulation, peaking at 25 minutes and declining thereafter. Analysis was done in at least 250 lysosomes in each time points. **(F)** Temporal correlation between mTORC1 recruitment and lysosome motility during nutrient restimulation derived from plot B and D. **(G–H)** Live-cell TIRF imaging of HAP-1 cells expressing endogenous RAPTOR-GFP (mTORC1) labelled with Lysotracker. Lysosomes positive for RAPTOR-GFP are more motile (**G**) and exhibit higher speeds (**H**) compared to RAPTOR-GFP negative lysosomes. The bar in all plots represents the mean, error bars represent SD; *P<0.05, **P<0.005, ****P<0.00005. The scale bar for all the images is 10 µm except for **G**, which is 2 µm.

To test whether mTORC1 recruitment caused lysosome motility, we turned to HAP1 cells expressing endogenously tagged GFP-RAPTOR (a core subunit of mTORC1 complex). Upon nutrient restimulation, RAPTOR-GFP colocalized with lysosomes in a time-dependent manner, peaking at 15 minutes, mirroring the trends observed in BS-C-1 cells (Figure S1B). Live-cell TIRF imaging of HAP1 cells in basal nutrient conditions revealed that lysosomes positive for mTORC1 (RAPTOR-GFP) are mobile, while mTORC1-negative lysosomes remained largely stationary (Figure 1G). Quantitative tracking over ∼1 minute at a 100-millisecond temporal resolution revealed that mTORC1-positive lysosomes moved at twice the speed of those lacking mTORC1 (Figure 1H). Long-term monitoring after nutrient restimulation (∼30 minutes) further confirmed that lysosomes became mobile upon acquiring mTORC1, with increased displacement over time (Figure S1C).

These findings demonstrate that mTORC1 recruitment is sufficient to trigger lysosomal motility, promoting their distribution across the cytoplasm. Notably, even at peak nutrient stimulation, only ∼55% of lysosomes colocalized with mTORC1 and exhibited enhanced motility. Both mTORC1 association and lysosomal motility declined following the peak, despite continued nutrient availability. These findings suggest that mTORC1-dependent lysosome motility is precisely regulated to control mTORC1 activity and maintain cellular homeostasis in response to nutrient cues.

### mTORC1 activation suppresses microtubule detyrosination

Microtubules are critical for directing long-range lysosomal transport (Guardia et al., 2016; Jordens et al., 2001; Korolchuk et al., 2011; Rocha et al., 2009; Rosa-Ferreira & Munro, 2011), and their post-translational modifications (PTMs) add a regulatory layer to this process (Konietzny et al., 2024). Notably, detyrosinated microtubules restrict lysosomal motility and promote fusion with autophagosomes (Mohan et al., 2019). We hypothesized that specific PTM-defined microtubule subsets may differentially regulate the motility of mTORC1-positive lysosomes, thereby influencing mTORC1 signalling activity.

To test this, we modulated mTORC1 activity in BS-C-1 cells using nutrient variability, genetic activation of mTOR, or rapamycin treatment, and quantified the levels of tyrosinated, detyrosinated, and acetylated tubulin — the major PTMs in epithelial cells. Structured illumination microscopy (SIM) analysis (see Methods, Figure. S2A) revealed that detyrosinated microtubules increased ∼1.5-fold under mTORC1-inhibiting conditions (starvation or rapamycin), with a concurrent decrease in tyrosinated microtubules (Figure 2A-B). Whereas high mTORC1 activity — induced by amino acid or serum stimulation, wild-type/hyperactive mTORC1, or RHEB overexpression — led to a ∼1.5-fold reduction in detyrosination (Figure 2A-B and S2B). These trends were validated by immunoblotting, where phospho-S6K1 and phospho-4EBP1 levels indicated mTORC1 activity (Figure 2C-G). In addition, we have observed same phenotype of reduction in detyrosinated microtubules on nutrient stimulation in another cell line, Osteosarcoma U2OS (Figure S2C-D) Notably, detyrosination levels inversely correlated with mTORC1 signaling (Figure 2A-G).

**Figure 2.**
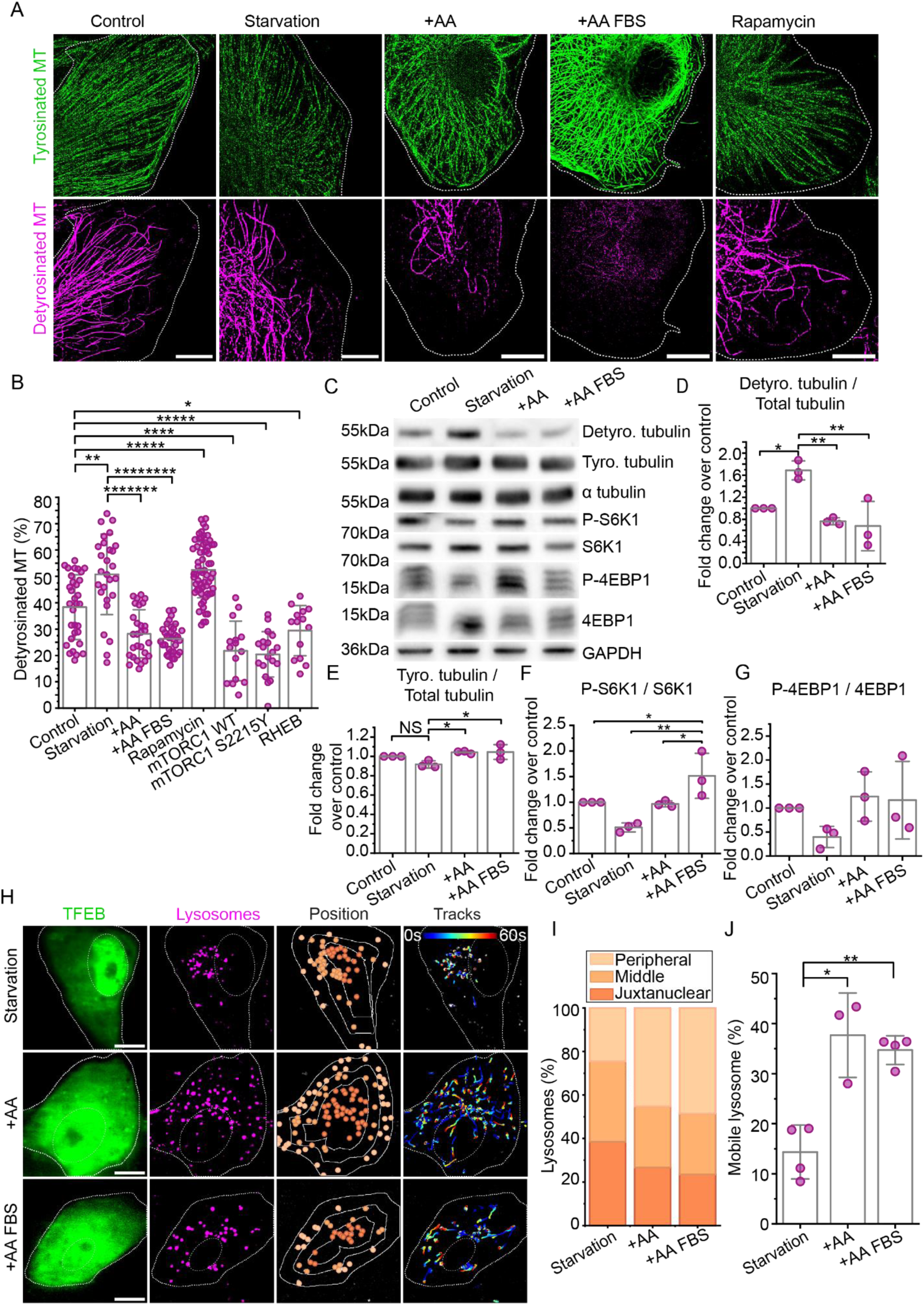
mTORC1 activation downregulate microtubule detyrosination. **(A)** Two colour SIM microscopy of tyrosinated (green) and detyrosinated (magenta) microtubule in BS-C-1 cells starved for 2 hr or starved and restimulated with AA or AA and FBS for 20 min or treated with rapamycin for 1hr. **(B)** Quantification of detyrosinated microtubules shows decrease in the percentage of detyrsoinated microtubule in AA or AA and FBS restimulation and increase in starvation or rapamycin treatment. **(C)**. Immunoblotting for tyrosinated and detyrosinated tubulin along with mTORC1 substrates S6K1 (phosphorylated and total) and 4EBP1 (phosphorylated and total) in BS-C-1 cells starved for 2 hr or starved and restimulated with AA or AA and FBS for 20 min **(D-G)** Densitometric quantifications of protein bands under different condition mentioned in **(C)** show the decrease in detyrosinated tubulin on mTORC1 activation represented by the phosphorylated level of S6K1 and 4EBP1.**(H)** Live-cell TIRF imaging and tracking of lysosomes (Lysotracker) in BS-C-1 cells transfected with EGFP TFEB under starvation and AA or AA and FBS restimulation conditions. Trajectories of individual lysosomes are shown with starting points in blue and endpoints in red.**(I)** Manual scoring of lysosomes positioned in juxtanuclear, middle, and peripheral zones of BS-C-1 cells as shown in **(H) (J)** Quantification of mobile lysosomes (displacement >1 µm) shows an increase in the mobile population in **(H)** with nutrient stimulation, where a total of 246, 370, and 420 lysosomes in starvation, AA, and AA and FB restimulation respectively were analysed from at least 3 cells per condition. The bar in all plots represents the mean, error bars represent SD; *P<0.05, **P<0.005, ****P<0.00005. The scale bar for all the images is 10 µm.

Acetylated microtubules in BS-C-1 cells are distributed into a larger proportion (55%) of acetylated-tyrosinated and a smaller proportion (45%) of acetylated-detyrosinated microtubules (Mohan et al., 2019). We observe that upon mTORC1 inhibition by starvation or rapamycin, the percentage of acetylated microtubules decreases, following the trend of tyrosination (Figure S2E-H). This suggests that, with mTORC1 suppression, the acetylated microtubule population that overlaps with tyrosination is reduced, while those that overlap with detyrosination remain.

Microtubule PTMs can modulate the dynamics of the microtubule (Wloga et al., 2017; Wloga & Gaertig, 2011), to check whether microtubule stability is modified under different nutrient conditions, we tracked the microtubule end binding protein (EB3), fused with EGFP in TIRF live cell imaging (Figure S2I). We found that under starvation the microtubule growth velocity is reduced by 0.026 µm/s compared to the control condition (Figure S2J).

To further connect changes in microtubule-PTMS with mTORC1 signalling and lysosomal dynamics, we examined lysosome positioning and motility under varying nutrient conditions using cells expressing GFP-tagged TFEB, a transcription factor (Roczniak-Ferguson et al., 2012) regulated by mTORC1 (Figure 2H-J). Under starvation, when mTORC1 is inactive and TFEB localizes to the nucleus (Roczniak-Ferguson et al., 2012; Settembre et al., 2011), lysosomes clustered in the perinuclear region and were largely immobile (Figure 2H-J). In contrast, nutrient-stimulated cells, showing cytoplasmic TFEB indicative of active mTORC1, exhibited dispersed, highly mobile lysosomes with increased anterograde movement (Figure 2H-J). Together, our results suggest that mTORC1 signalling promotes microtubule-tyrosination and microtubule dynamics, which likely facilitates the rapid distribution of lysosomes positive for mTORC1 by enhancing their long-range transport.

### mTORC1-positive lysosomes preferentially traffic along tyrosinated microtubules for rapid cytoplasmic distribution

Next, we investigated whether tyrosinated microtubules promote the distribution and long-range transport of lysosomes that carry mTORC1. To address this, we first quantified the fraction of lysosomes associated with mTORC1 using two-color TIRF imaging of BS-C-1 cells under basal conditions and following amino acid stimulation. We then employed two-color super-resolution STORM imaging to precisely resolve the localization of mTORC1-positive lysosomes relative to tyrosinated microtubules. Strikingly, mTORC1-positive lysosomes showed a strong preference for tyrosinated microtubules. Under basal conditions, 66.4 ± 10.1% of mTORC1-positive lysosomes were localized on tyrosinated microtubules, increasing to 81.8 ± 11.8% with amino acid stimulation (Figure 3A-B). This enrichment was corroborated by two-color SIM imaging, where 67 ± 5.6% of total mTORC1 puncta localized on to tyrosinated microtubules under control conditions, and increasing to 79.2 ± 7.7% after stimulation (Figure S3A&B). By contrast, less than 5% of mTORC1 puncta localized to detyrosinated microtubules in either condition. Together, these data demonstrate that mTORC1-positive lysosomes predominantly associate with tyrosinated microtubules, particularly under nutrient-rich conditions.

**Figure 3.**
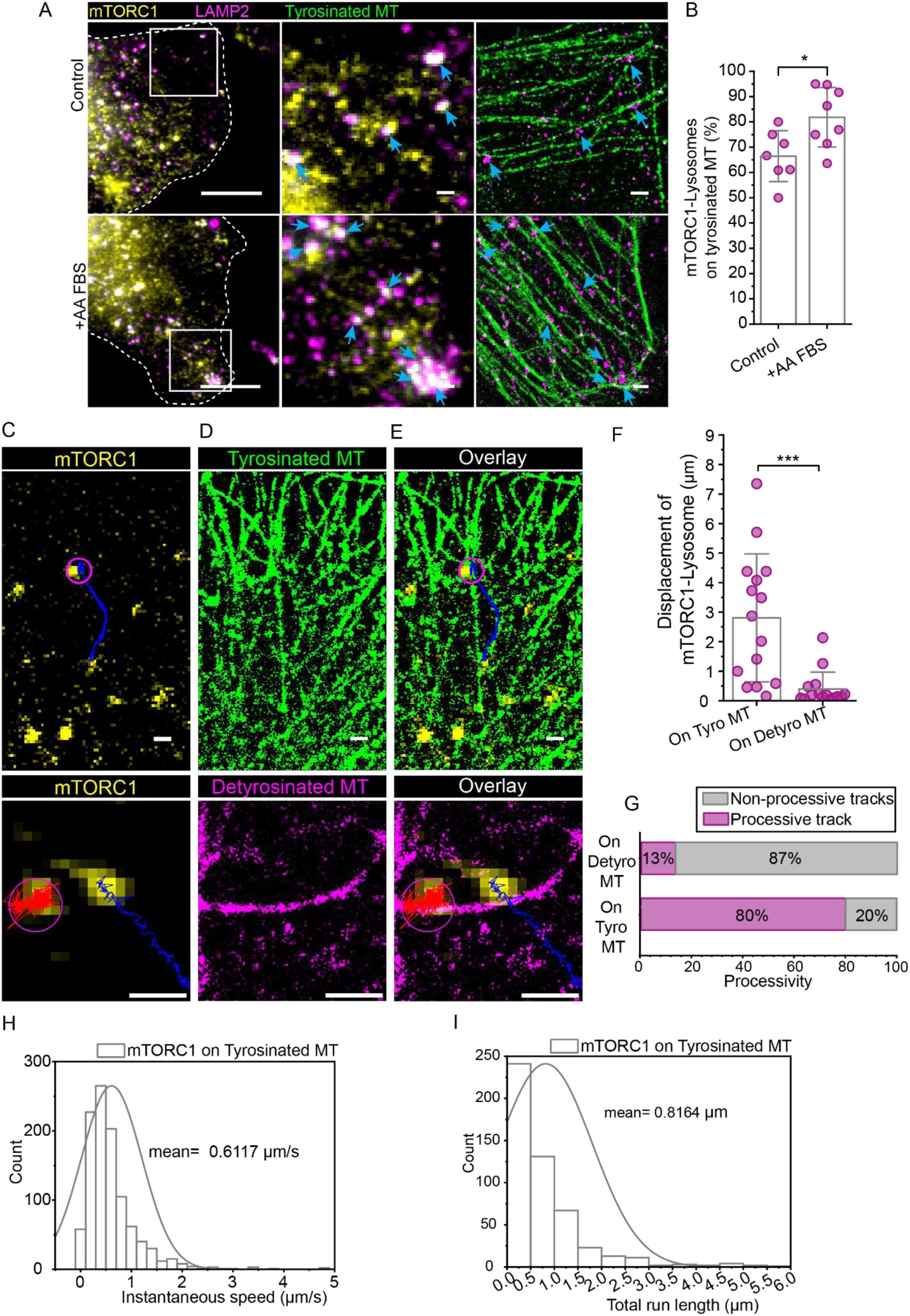
mTORC1-associated lysosomes localise and moves on tyrosinated MT. **(A)** Two-colour STORM microscopy of tyrosinated (green) microtubule and lysosome (magenta) microtubule and conventional endogenous mTORC1 (yellow) in BS-C-1 cells untreated or starved and restimulated with AA and FBS for 20 min. **(B)** Manual scoring of mTORC1-positive lysosomes on tyrosinated microtubules shows an increase in the percentage of mTORC1-positive lysosomes localised to tyrosinated microtubules on AA FBS restimulation. Correlative live-cell and STORM, **(C)** shows the tracking of mTORC1 YFP (yellow), which is aligned on the **(D)** STORM image of tyrosinated microtubule (green) or detyrosinated microtubule (magenta) and overlayed **(E)**, track of mTORC1 on tyrosinated microtubule is represented in blue while that on the detyrosinated microtubule is represented in red colour**. (F-G)** Tracking analysis of mTORC1-positive lysosomes on tyrosinated or detyrosinated microtubules, **(F)** displacement and **(G)** processivity of the tracks, shows that the mTORC1-lysosome is having reduced displacement and hindered motility on the detyrosinated microtubules. **(H)** Instantaneous speed and **(I)** total run length of mTORC1 on tyrosinated microtubule. A total of 15 mTORC1-positive lysosomes on tyrosinated or detyrsoinated microtubule were analysed from two cells. The bar in the plot represents the mean, error bars represent SD; *P<0.05, **P<0.005, ****P<0.00005. The scale bar for **(A)** the images is 10 µm and 1 µm for inset zoom images. The scale bar for **(C-E)** is 1 µm.

To test whether tyrosinated microtubules actively support the long-range transport of mTORC1-positive lysosomes, we performed correlative live-cell and super-resolution STORM imaging. Firstly, we tracked individual mTORC1 puncta in live BS-C-1 cells using TIRF microscopy for ∼1 minute at 100-millisecond intervals. Cells were then fixed and immunostained for either tyrosinated or detyrosinated tubulin, followed by STORM imaging of the microtubules. Overlaying the trajectories onto the high-resolution microtubule maps revealed that mTORC1 puncta moved preferentially on tyrosinated microtubules (Figure 3 C-E).

Single-particle tracking analysis confirmed that mTORC1 moved with significantly greater (6 folds) displacement (Figure 3F and G) compared to detyrosinated ones. Since all the tracks of mTORC1 on detyrosinated microtubule are non-processive, we have calculated speed (Figure 3H) and run lengths (Figure 3I) of mTORC1 on tyrosinated microtubules only. These findings establish that tyrosinated microtubules serve as specialized tracks for the efficient and sustained transport of mTORC1-positive lysosomes to distribute them across the cytoplasm.

### mTORC1 downregulates microtubule detyrosination through S6K1

Once validated that the mTORC1 upregulates microtubule tyrosination to enhance its activity by promoting the rapid peripheral movement and positioning of lysosomes that are positive for mTORC1 throughout the cytoplasm, we investigated to find the molecular player responsible for the regulation of microtubule tyrosination or detyrosination mediated by mTORC1. To this end, we screened the kinase in the mTORC1 signal transduction, upstream and downstream of mTORC1 (AKT, mTOR, S6K1), for possible interaction and regulation of microtubule tyrosination/detyrosination enzymes (Tubulin Tyrosine Ligase and Vasohibin) using the PhosphoSite Plus tool (Hornbeck et al., 2015). We found that mTORC1 substrate S6K1 can have a phosphorylating site on VASH2 at serine 332, which is conserved throughout the mammalian species (Figure S4A). To validate that mTORC1 regulates microtubule detyrosination through S6K1, we found a drug that specifically inhibits S6K1 activity without affecting its phosphorylation, PF-4708671 (Rosner et al., 2012). We modulated the activity of S6K1 by treating with DMSO or PF-4708671 (2 µM) (IC50: 0.16 μM, (Pearce et al., 2010)) under varying nutrient conditions. Structured illumination microscopy (SIM) analysis revealed that AA (Amino Acid) restimulation-induced reduction of detyrosinated microtubules is blocked by the S6K1 inhibitor (PF-4708671) (43.33±20.38%) (Figure 4A-B). These trends were validated by immunoblotting, where the level of detyrosinated microtubule is increased significantly upon S6K1 inhibitor treatment regardless of the nutrient status of the cell (Figure 4C-D), and the mTORC1 activity is also reduced (Figure 4C&G). As reported by Pearce, et al., (2010), we also see increased (but not significant) phosphorylation of S6K1 in the presence of PF-4708671 (Figure 4C&F) even in the absence of growth factor. The mTORC1 activity is significantly reduced in the amino acid restimulation condition in the presence of the S6K1 inhibitor compared to that of the control, the reason is possibly because of feedback inhibition of IRS-1 by phosphorylated S6K1 in the high AA condition, thus in turn reducing the mTORC1 activity (Tremblay et al., 2007). These results indicate that the S6K1 is downregulating the level of detyrosinated microtubule, acting downstream of mTORC1 upon activation.

**Figure 4.**
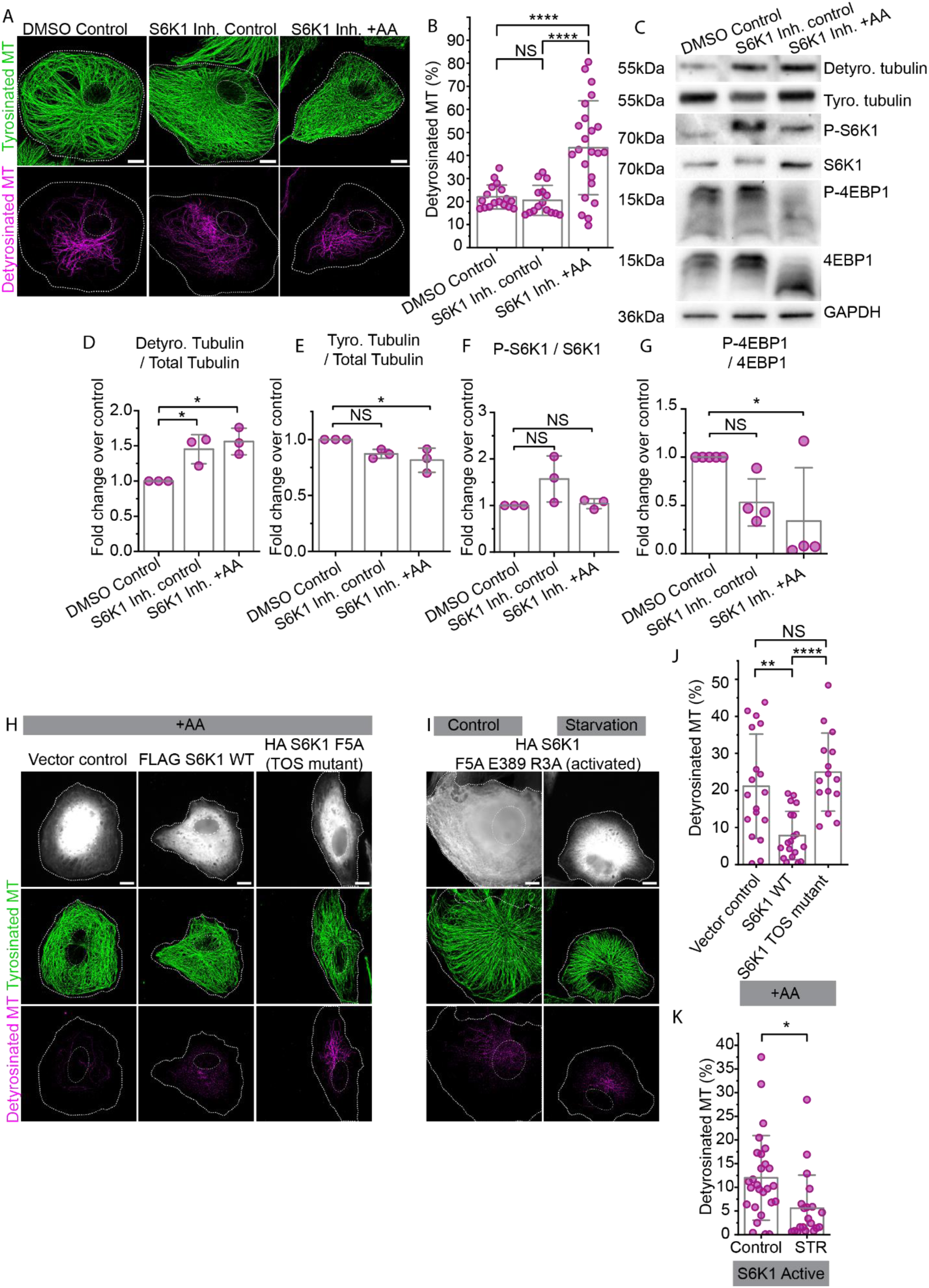
mTORC1 downregulates microtubule detyrosination through S6K1. **(A)** Two-colour SIM microscopy of tyrosinated (green) and detyrosinated (magenta) microtubules in BS-C-1 cells untreated in the presence of DMSO or untreated or starved for 2 hr and restimulated with AA for 20 min in the presence of S6K1 inhibitor (PF-4708671) (2 µM). **(B)** Quantification of detyrosinated microtubules shows an increase in the detyrosinated microtubule on AA restimulated in the presence of the S6K1 inhibitor. **(C)**. Immunoblotting for tyrosinated and detyrosinated tubulin along with mTORC1 substrates S6K1 (phosphorylated and total) and 4EBP1 (phosphorylated and total) in BS-C-1 cells treated as in **(A). (D-G)** Densitometric quantifications of protein bands under different conditions mentioned in **(C),** show the increase in detyrosinated tubulin in the presence of S6K1 inhibitor in untreated and AA restimulation conditions. **(H and I)** Two-colour SIM microscopy was performed on tyrosinated (green) and detyrosinated (magenta) microtubules in BS-C-1 cells that overexpress either the vector backbone or FLAG S6K1 WT or HA S6K1 F5A (TOS mutant), or HA S6K1 F5A E389 R3A (activated). The cells were either untreated, starved for 2 hours, or starved and then restimulated with AA for 20 minutes. **(J and K)** Quantification of detyrosinated microtubules shows inhibition in the reduction of detyrosinated microtubules on AA restimulation in the cells overexpressed with TOS mutant S6K1. The bar in all plots represents the mean, and error bars represent SD; *P<0.05, **P<0.005, ****P<0.00005. The scale bar for all the images is 10 µm.

To further validate this phenotype by genetical modulation of S6K1, we obtained the S6K1 TOS mutant (F5A), which impairs recognition and activation by mTORC1, and the S6K1 kinase-activated variant. SIM analysis of cells transfected with S6K1 WT or S6K1 TOS mutant under AA restimulation revealed that WT overexpression further reduced the detyrosinated microtubule level to 7.84 ± 6.5%, whereas TOS mutant overexpression did not significantly reduce the level of detyrosinated microtubule (24.92 ± 10.54%) compared to the vector control (21.15 ± 14.07%) (Figure 4 H&J). Concurrently, cells transfected with the kinase-activated variant of S6K1 reduced the level of detyrosinated microtubule to 11.99 ± 8.93% in the basal condition and blocked the increase of detyrosinated microtubule even in nutrient starvation (5.6 ± 6.98) (Figure 4 I&K). All these results confirm that S6K1 is downregulating the level of detyrosinated microtubule, acting downstream of mTORC1 upon activation.

### Microtubule tyrosination promotes mTORC1 activity

Given our observations that nutrient stimulation decreases microtubule detyrosination and that tyrosinated microtubules enable the rapid trafficking of mTORC1-containing lysosomes, we next aim to investigate whether the tyrosination/detyrosination state of microtubules directly regulates lysosomal redistribution and mTORC1 activity. To this end, we overexpressed tubulin tyrosine ligase (TTL) — TTL mscarlet, or Vasohibin (VASH) — VASH2 mCherry, the enzymes known for catalyzing tyrosination and detyrosination of microtubules, respectively, in BS-C-1 cells and validated their activity by imaging the modulation of the microtubule-PTMs levels (Figure S5A).

Notably, cells overexpressing VASH, (high levels of detyrosinated microtubules) exhibited ∼1.5-fold increase in lysosomal confinement to the perinuclear region compared to controls (Figure 5A-B). Congruently, there was a ∼3.7-fold reduction in the total mobile lysosome population and a roughly 1.6-fold decrease in anterogradely moving lysosomes in cells with increased detyrosinated microtubules compared to controls (Figure 5A-D). In contrast, TTL overexpression (high levels of tyrosinated microtubules) caused lysosomes to become more dispersed and mobile, although this difference was not statistically significant from control conditions, likely because TTL acts on tubulin monomers rather than on polymerized microtubules, thereby allowing detyrosinated microtubules to persist despite TTL overexpression.

**Figure 5.**
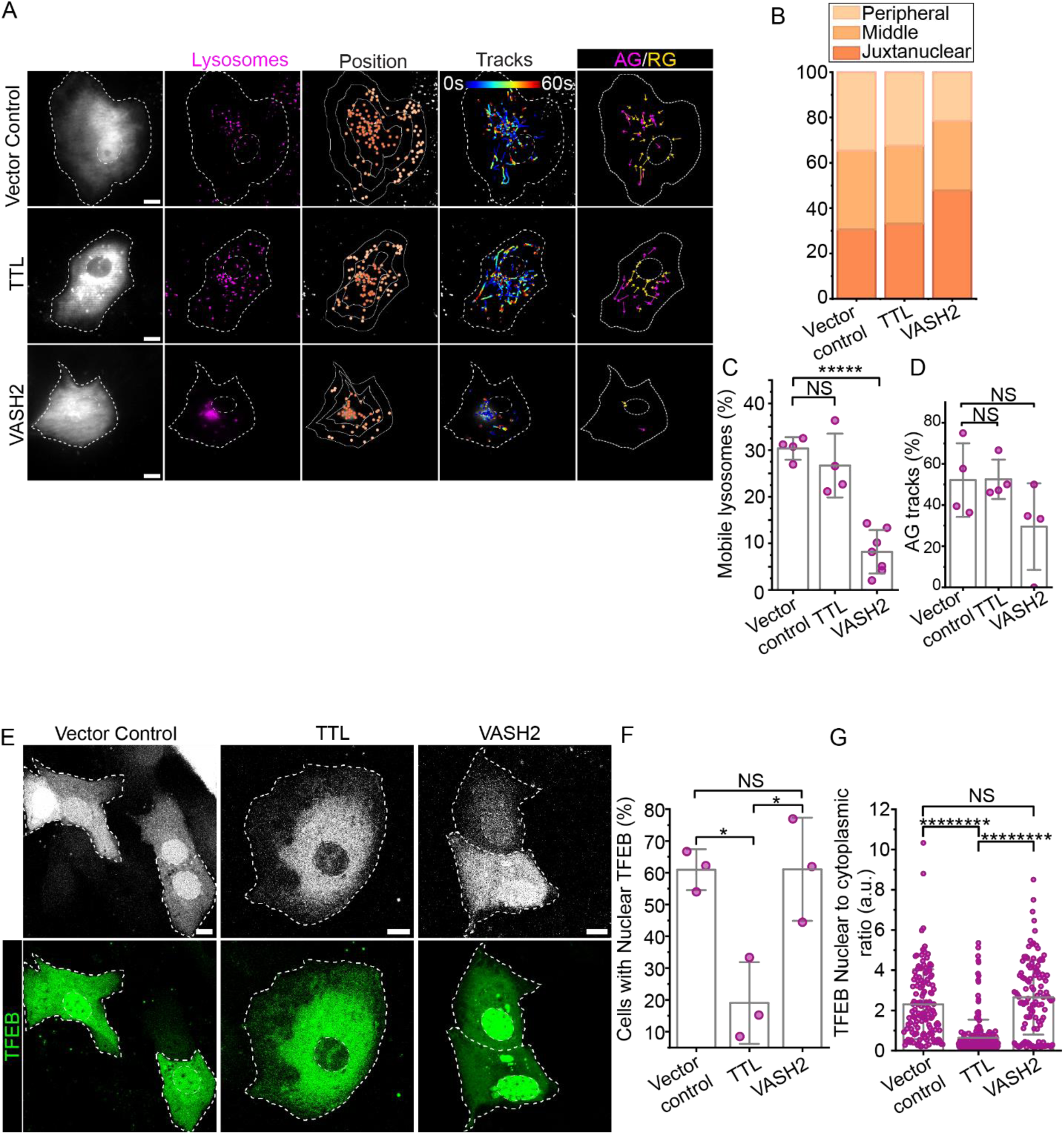
mTORC1 activity requires microtubule tyrosination. **(A)** Live-cell TIRF imaging and tracking of lysosomes (Lysotracker) in BS-C-1 cells transfected with vector control or TTL mscarlet or VASH2 mCherry. Trajectories of individual lysosomes are shown with starting points in blue and endpoints in red. Tracks are categorized into anterograde (AG, magenta) and retrograde (RG, yellow) movement**. (B)** Manual scoring of lysosomes positioned in juxtanuclear, middle, and peripheral zones of BS-C-1 cells in **(A).** Quantification of **(C)** mobile lysosomes (displacement >1 µm) and **(D)** anterograde movement shows a decrease in the mobile population and decrease in anterograde movement for BS-C-1 cells in **(A)** under VASH2 overexpression. A total of 512, 510, and 1084 lysosomes in vector control, TTL mscarlet, and VASH2 mCherry overexpression conditions, respectively were analyzed from at least 4 cells per condition. **(E)** BS-C-1 cells transfected with EGFP TFEB along with vector control or TTL mscarlet or VASH2 mCherry. **(F)** Manual quantification of cells with nuclear TFEB shows that the percentage of cells with nuclear TFEB reduced on TTL mscarlet overexpression condition, where a total of 98, 111, and 43 cells in vector control, TTL mscarlet, and VASH2 mCherry overexpression conditions, respectively were analysed from 3 replicates per condition. **(G)** Intensity based quantification of TFEB in unit nuclear to cytoplasmic area reveals that TFEB nuclear-to-cytoplasmic ratio decreases on overexpression with TTL mscarlet. The bar in all plots represents the mean, error bars represent SD; *P<0.05, **P<0.005, ****P<0.00005. The scale bar for all the images is 10 µm.

Next, we assessed the impact of changes in tyrosination/detyrosination levels on mTORC1 activity by observing the subcellular localization of TFEB. TTL overexpression resulted in a ∼3-fold reduction in the proportion of cells with nuclear TFEB and a ∼2.7-fold decrease in the nuclear-to-cytoplasmic ratio of the EGFP TFEB signal, compared to controls (Figure 5E-G), indicating significantly increased mTORC1 activity. Conversely, VASH overexpression showed a modest increase in the nuclear TFEB signal, compared to the control condition (Figure 5E-G), however, this was non-significant when compared to the control condition, likely because EGFP TFEB overexpression already saturated the proportion of cells with nuclear TFEB, which could not be further increased by VASH overexpression. Together, our results demonstrate that the tyrosination status of microtubules promotes mTORC1 activity and either suppressing detyrosination or upregulating tyrosination can enhance mTORC1 signaling in epithelial cells.

### Restoring microtubule detyrosination reverses mTORC1 hyperactivity and cancer cell migration

Aberrant mTORC1 activity is a hallmark of many cancer types. mTORC1 hyperactivity promotes uncontrolled cell growth, proliferation, and metastasis. Our findings demonstrate that microtubule tyrosination is a key regulator of mTORC1 activity. Therefore, we hypothesised that in cancer types where mTORC1 hyperactivity is associated with increased microtubule tyrosination, the cancer phenotypes could be reversed by restoring mTORC1 activity to normal levels through downregulating tubulin tyrosination.

Using differential gene expression data from the UALCAN database (Chandrashekar et al., 2022), we examined the relative expression of TTL, VASH1/2, and mTOR between normal and tumour samples across multiple cancer types, where we narrowed our focus on prostate adenocarcinoma, which showed a positive correlation of mTOR with TTL and a negative correlation with VASH2 and was considered for further investigation (Figure S6A). We used prostate carcinoma cell line models 22RV1, and LNCaP to measure the correlation between mTOR and microtubule tyrosination/detyrosination levels and compared their expression levels to a benign prostate carcinoma cell line PNT2. We observed that in prostate carcinoma cell line models 22RV1 and LNCaP, the tyrosinated microtubule level was upregulated and detyrosinated microtubules levels were downregulated along with upregulated mTORC1 activity reflected by higher phosphorylated S6K1 levels (Figure 6 A&B, S6 B-D), in comparison to a benign prostate carcinoma cell line PNT2. These correlations were consistent in an endometrial adenocarcinoma cell line JHUEM7 (Figure 6 A&B, S6 B-D), which is a well-established model to harbour mTOR mutation (S2215Y) leading to its hyperactivity (Grabiner et al., 2014).

**Figure 6.**
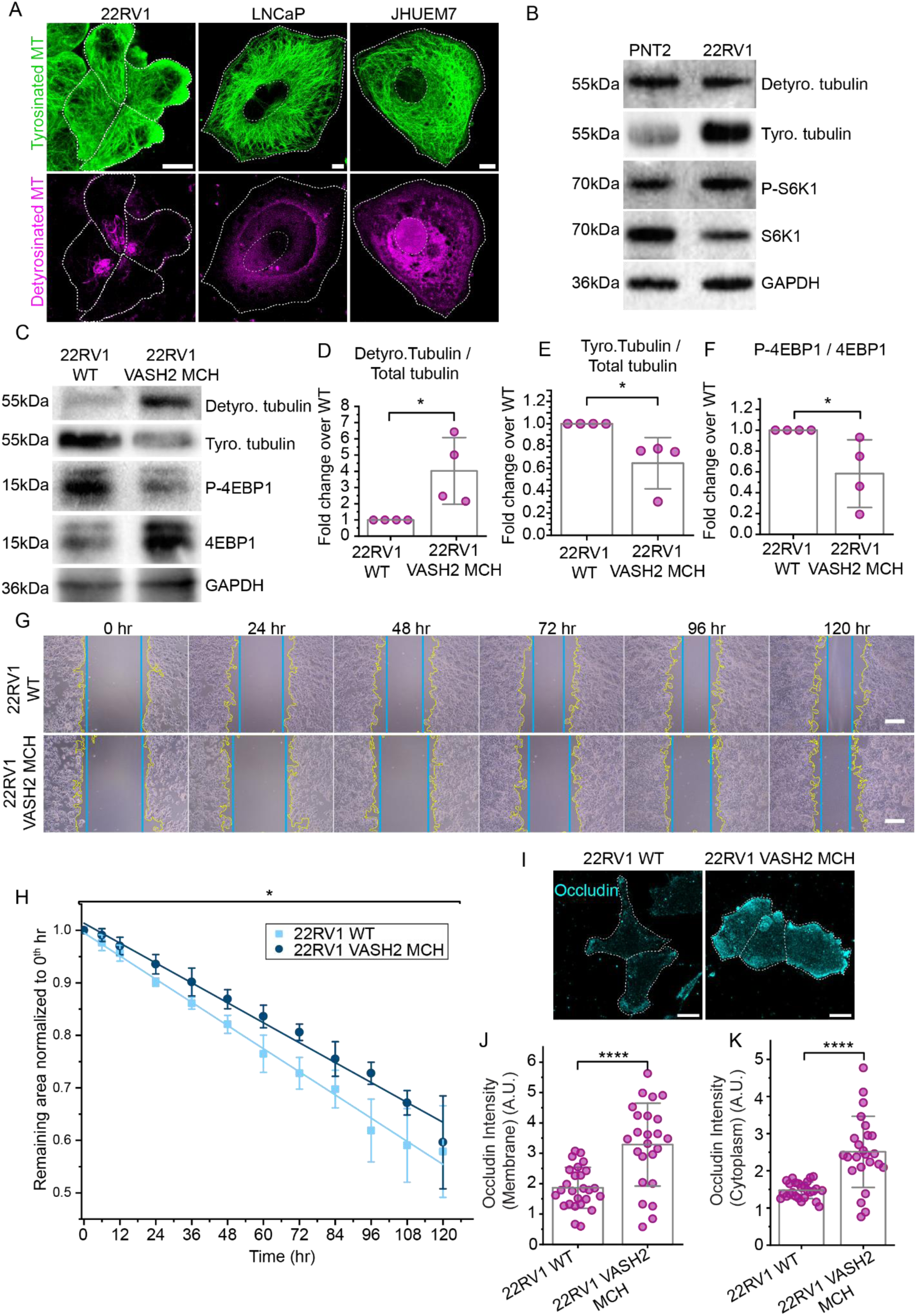
Cancer phenotypes with aberrant mTORC1 activity can be rescued by downregulating microtubule tyrosination. **(A)** Confocal imaging of tyrosinated (green) and detyrosinated (magenta) microtubules showing abundance of tyrosinated microtubule in 22RV1, LNCaP and JHUEM7 cells. **(B)** Immunoblotting for tyrosinated and detyrosinated tubulin along with mTORC1 substrate S6K1 (phosphorylated and total) in PNT2 and 22RV1 cells showing the mTORC1 activity and higher level of tyrosinated microtubules in 22RV1 cells. **(C)** Immunoblotting for tyrosinated and detyrosinated tubulin along with mTORC1 substrate 4EBP1 (phosphorylated and total) in WT and VASH2 mCherry stably overexpressed 22RV1 cells **(D-F)** Densitometric quantifications of protein bands in **(C)** show the decrease in tyrosinated tubulin and mTORC1 activity represented by the phosphorylated level of 4EBP1 in VASH2 mCherry stably overexpressed 22RV1 cells. **(G)** Wound healing assay in WT 22RV1 and VASH2 mCherry stably overexpressed 22RV1 showing the wound closure for 5 days. **(H)** Quantification of wound closure area showing delayed wound closure and decreased migration rate **(Figure S6B)** in VASH2 mCherry stably overexpressed 22RV1 cells. **(I)** Confocal microscopy of Occludin in WT and VASH2 mCherry stably overexpressed 22RV1 cells. **(J&K)** Intensity based quantification of Occludin in membrane as well as cytoplasmic region of the cell, reveals that the expression of occludin is increased on VASH2 mCherry stably overexpressed 22RV1 cells. The bar in all plots represents the mean, error bars represent SD; *P<0.05, **P<0.005, ****P<0.00005. The scale bar for all the images is 10 µm.

Among all the cell lines, 22RV1 showed the highest mTORC1 hyperactivity, reflected by elevated levels of P-S6K1. Hence, to test whether modulating tyrosination/detyrosination levels could revert the mTORC1 levels to normal, we generated a 22RV1 cell line stably expressing VASH2 to upregulate microtubule detyrosination in these cells. Notably, VASH2 expression reduced the level of tyrosinated microtubules and the phosphorylation levels of 4EBP1, the downstream substrate of mTORC1 (Figure 6C-F). Consistently, transient knockdown of the TTL enzyme, leading to downregulation of microtubule tyrosination, resulted in a reduction in mTORC1 activity in both 22RV1 and LNCAP cells (Figure S6 E-L). Furthermore, VASH2 overexpression in 22RV1 cells resulted in reduced cell migration (Figure 6 G&H, S6 M).Moreover, we checked the expression level of tight junction integral protein occludin (whose expression is shown to be downregulated in multiple cancer types (Büyücek et al., 2025; Osanai et al., 2006)) in WT and VASH2 stably expressing 22RV1 cells (Figure 6I). Fluorescent Intensity based quantification of occludin in cytoplasmic as wells as membranal region revealed ∼1.7-fold increase of occludin expression in VASH2 expressed cells (Figure 6 J&K), this indicates that tumorigenic property of 22RV1 cells is reduced in the cells with higher microtubule detyrosination. Together, these findings suggest that the hyperactivity of mTORC1 can be reversed by downregulating microtubule tyrosination or by upregulating detyrosination, thereby preventing uncontrolled cancer cell proliferation and migration.

## Discussion

We identify a direct coupling between lysosomal positioning, microtubule PTMs, and nutrient-dependent mTORC1 activity. Our findings establish that only the subset of lysosomes decorated with mTORC1 display enhanced motility and peripheral positioning under nutrient stimulation, while others remain largely immobile. This selectivity suggests a feedback mechanism that ensures nutrient homeostasis by dynamically regulating the fraction of motile, mTORC1-positive lysosomes. A central outcome of our study is the demonstration that tyrosinated and detyrosinated microtubules serve distinct and complementary roles in lysosome biology: tyrosinated microtubules facilitate mTORC1 activity, whereas detyrosinated microtubules promote autophagy. This partitioning provides a mechanistic basis for how nutrient availability is translated into opposing lysosomal outputs. Under nutrient-rich conditions, lysosomes containing mTORC1 preferentially move along tyrosinated microtubules to the cell periphery, enhancing signaling. By contrast, during starvation, detyrosinated microtubules accumulate and support autophagosome–lysosome fusion, promoting catabolic function. Thus, the tight regulation of microtubule tyrosination/detyrosination status controls lysosome motility and thereby maintains nutrient homeostasis.

Our data refine earlier reports on nutrient-dependent changes in microtubule PTMs. Previous work in HeLa cells suggested that starvation stabilizes microtubules through acetylation, thereby supporting autophagy (Geeraert et al., 2010). In BS-C-1 cells, however, we observed increased detyrosination but decreased acetylation upon starvation. Such discrepancies are likely cell- and tissue-specific, reflecting the differential regulation of αTAT1 and HDAC6, enzymes that control microtubule acetylation(Bhattacharya et al., 2025). Indeed, acetylation has been reported to increase under high nutrient conditions (Parker et al., 2019), and cancer studies have also highlighted context-dependent acetylation dynamics. Our findings suggest that in BS-C-1 cells, acetylation interacts with both tyrosinated and detyrosinated microtubule subsets, thereby shifting the balance between acetylated tyrosinated and non-acetylated detyrosinated populations in response to nutrient status.

Manipulating tyrosination levels revealed additional layers of regulation. Increasing tyrosinated tubulin enhanced mTORC1 signalling but only modestly boosted lysosomal motility (only 10% variation, Figure 5A-D), suggesting that tyrosinated microtubules may regulate mTORC1 through mechanisms beyond transport. One candidate is CLIP-170, a CAP-Gly protein that requires tyrosinated tubulin for microtubule binding (Bieling et al., 2008; Peris et al., 2006). mTORC1 can phosphorylate CLIP-170 (Choi et al., 2002), and CLIP-170 has been shown to capture cargo at microtubule plus-ends (Lomakin et al., 2009). Together, these findings suggest that tyrosinated microtubules may provide a signalling platform where mTORC1 stabilizes lysosomal residence at the periphery via CLIP-170.

Our results also integrate with emerging evidence that lipid composition and motor recruitment define lysosomal behaviour. Lysosomal PI4P–PI3P switches influence mTORC1 activity (Ebner et al., 2023), while distinct kinesins engage with nutrient-state-specific microtubules. KIF1B, KIF18B, and KIF16B preferentially transport lysosomes on tyrosinated tracks, whereas KIF5B can operate on acetylated or detyrosinated microtubules (Guardia et al., 2016; Korolchuk et al., 2011; Li et al., 2023). These mechanisms converge with our observation that mTORC1-positive lysosomes selectively engage with tyrosinated microtubules, reinforcing the idea that lipid cues, kinesin identity, and microtubule PTMs act in concert to coordinate lysosomal nutrient responses.

Beyond nutrient signalling, our findings have implications for cancer biology. Dysregulation of microtubule PTMs is a recurring theme in malignancy: acetylation supports migration and chemoresistance in lung cancer (Wattanathamsan et al., 2021), while detyrosination promotes EMT and metastasis in breast and ovarian cancers (Koyanagi et al., 2021; Whipple et al., 2010). Tyrosination, conversely, is linked to colorectal cancer progression and paclitaxel resistance (Banerjee, 2002; Yao et al., 2017). These conflicting phenotypes likely reflect context-dependent regulation of PTMs across tissues. By directly connecting tyrosinated microtubules to mTORC1 activity, we reveal a mechanism that could reconcile some of these inconsistencies. Specifically, our data suggest that reducing tyrosination may attenuate hyperactive mTORC1 signalling and cancer cell migration, highlighting tyrosination as a potential therapeutic target to mitigate aberrant mTORC1 signalling mediated malignancies.

Several clinically approved drugs already exploit microtubule stabilization (e.g., paclitaxel, docetaxel, eribulin). Targeting detyrosination has also been explored: parthenolide suppresses metastatic traits in breast and colorectal cancers (Liu et al., 2017; Whipple et al., 2013), while EpoY enhances chemosensitivity in lung cancer by disrupting detyrosinated microtubule–KIF3C interactions (J. Wang et al., 2024). Our findings extend this therapeutic landscape by nominating tyrosination as a novel, context-specific target, particularly relevant to cancers driven by aberrant mTORC1 signaling.

In conclusion, we propose a unifying model in which nutrient availability remodels the microtubule cytoskeleton to partition lysosomal function: tyrosinated microtubules enable mTORC1 activity and anabolic signalling, while detyrosinated microtubules promote autophagy. This framework integrates lysosomal lipid remodelling, kinesin recruitment, and microtubule PTM dynamics into a coherent regulatory axis. Future studies will be required to elucidate how different cell types fine-tune this system, how feedback between mTORC1 and microtubule modifiers operates, and whether targeted manipulation of tyrosination can provide therapeutic benefits in cancer and metabolic diseases.

## Materials and Methods

### Cell culture

African green monkey (*Chlorocebus pygerythrus*) kidney cells (BS-C-1, CCL-26; American Type Culture Collection) were grown in complete growth media (CGM) (Minimum essential medium with nonessential amino acids containing 10% [vol/vol] FBS, 2 mM L-glutamine, 1 mM sodium pyruvate, and penicillin-streptomycin) at 37°C and 5% CO₂.

A stable transfected cell line for mTOR YFP was derived from the BS-C-1 line by transfection with the peYFP-C1-mTOR plasmid (Addgene, 73384), and clone selection was done in CGM with 600 µg/ml Geneticin (G418 Sulfate, 10131035; In-vitrogen).

The CRISPR Cas9-edited Haploid cell (Hap1) containing Raptor GFP (a kind gift from Prof. Nicholas T Ktistakis) were grown in complete growth media (Iscove’s Modified Dulbecco’s Medium with 10% [vol/vol] FBS and Antibiotic-Antimycotic) at 37°C and 5% CO₂.

Cancer cell lines PNT2, 22RV1, and LNCaP (kind gift from Prof. Bushra Ateeq) were grown in complete growth media (RPMI 1640 Medium, GlutaMAX™ Supplement containing 10% [vol/vol] FBS, and 0.5% penicillin-streptomycin) at 37°C and 5% CO₂.

### Lentiviral stable clone generation

HEK293T cells were transfected with pTRIP-CAG-mSVBP-mVASH2_X1-mCherry along with the packaging plasmids psPAX2 and pCMV-VSV-G at the ratio of 5:3:2. Two days after transfection, the supernatant containing lentiviruses was collected. Collected lentiviral particles were used to infect 22RV1 cells. 48 hr after infection, the cells were treated with selection media containing puromycin (1-5 µg/ml) for one week to select knockout cells. The knockouts were confirmed by western blotting.

### Nutrient and drug Treatment

For starvation of BS-C-1 cells, they were treated with starvation media (Krebs-Ringer’s solution (pH 7.4) (KRBH), supplemented with 4.5 mM glucose, 0.1% bovine serum albumin, and 1 mM sodium pyruvate) for 2 hr (Dyachok et al., 2016). For restimulation, BS-C-1 cells were starved and re-simulated with amino acid recovery media (DMEM without amino acid and low glucose containing 2X MEM amino acid in one-letter code R, C, H, I, L, K, M, F, T, W, Y, V from Invitrogen) with or without 10% (vol/vol) FBS as per requirement for 15-20 min or for the indicated time.

For starvation of Hap1 Raptor GFP cells, cells were treated with starvation media (140 mM NaCl, 1 mM CaCl₂, 1 mM MgCl₂, 5 mM glucose, 20 mM Hepes, 5 mM KCl, pH 7.4 containing 1% BSA) for 1 hour (Manifava et al., 2016). For restimulation cells were starved and re-simulated with amino-acid recovery media (DMEM without amino acid and low glucose containing 2X MEM amino acid in one-letter code R, C, H, I, L, K, M, F, T, W, Y, and V from Invitrogen) for 15 min.

For starvation of U2 OS cells, cells were starved with Hank’s Balanced Salt Solution (HBSS) for 2 hr (Shang et al., 2011)and recovered with amino-acid recovery media (HBSS containing 10X MEM amino acid in one letter code R, C, H, I, L, K, M, F, T, W, Y, and V from Invitrogen) for 30 min.

For rapamycin treatment, BS-C-1 cells were treated with 0.2 µm of rapamycin (Merck, R0395) or DMSO for 1 hr (Singh et al., 2013). For S6K1 inhibition, BS-C-1 cells were incubated in 2 µm of PF-4708671 (Merck, 559273) or DMSO. For restimulation with the S6K1 inhibitor, PF-4708671 was added to the starvation media after 2 hr of starvation and incubated for 1 hr, followed by restimulation in the presence of PF-4708671(Pearce et al., 2010; Rosner et al., 2012).

### Plasmids and Transfection

Cells were seeded in 8-well chambered cover glass or confocal disc 12-24 hr before the transfection and maintained in CGM. After 24 hr, transfection was performed using the Roche Effectene transfection kit with the manufacturer’s protocol. For transfection of mTORC1 plasmids, BS-C-1 cells were transfected with Raptor GFP, myc mTOR, and mlst8 at a ratio of 1:2:2 using the above-mentioned reagents.

### Key resource table

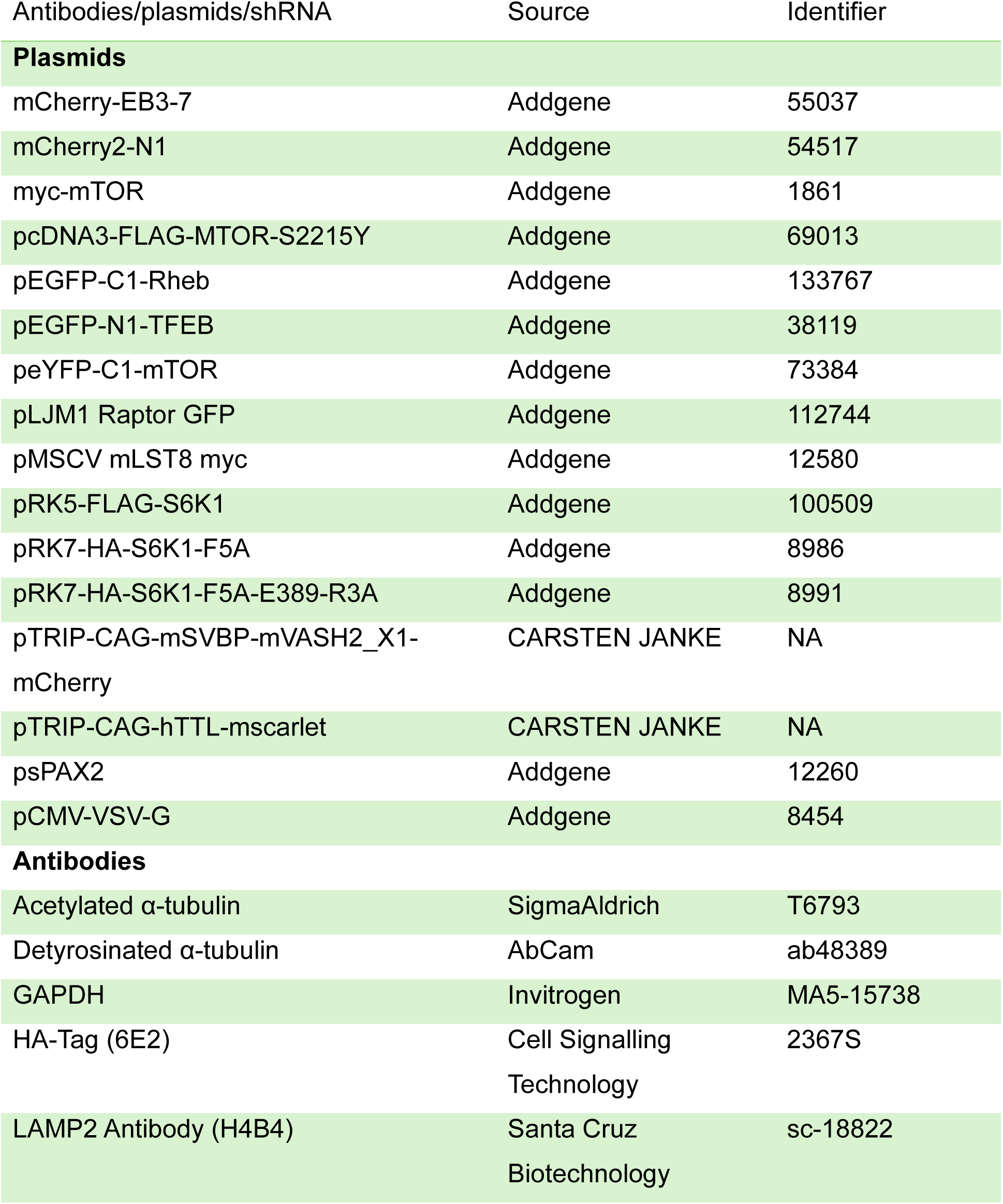
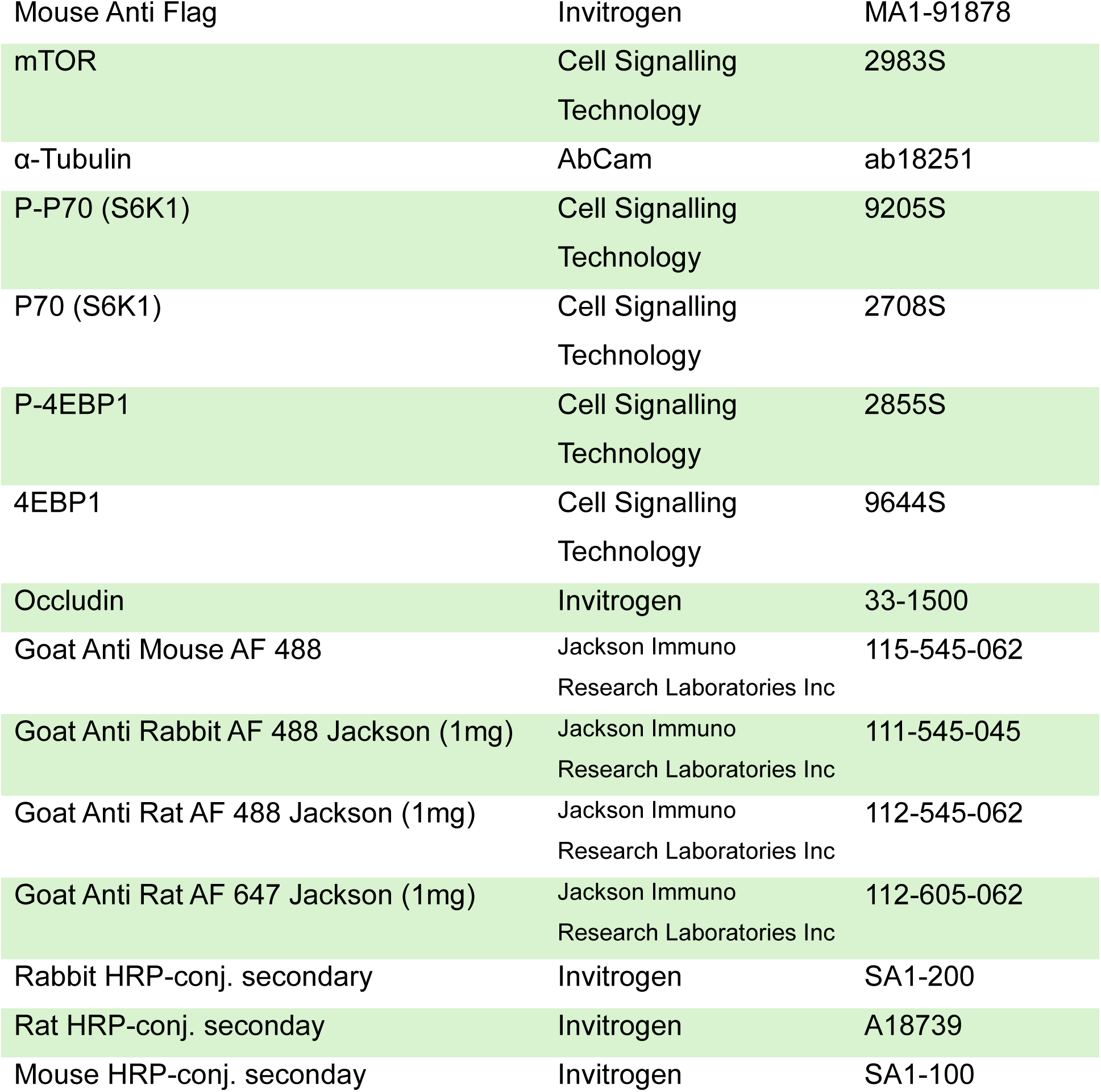

| Antibodies/plasmids/shRNA | Source | Identifier |
| --- | --- | --- |
| **Plasmids** |  |  |
| mCherry-EB3-7 | Addgene | 55037 |
| mCherry2-N1 | Addgene | 54517 |
| myc-mTOR | Addgene | 1861 |
| pcDNA3-FLAG-MTOR-S2215Y | Addgene | 69013 |
| pEGFP-C1-Rheb | Addgene | 133767 |
| pEGFP-N1-TFEB | Addgene | 38119 |
| peYFP-C1-mTOR | Addgene | 73384 |
| pLJM1 Raptor GFP | Addgene | 112744 |
| pMSCV mLST8 myc | Addgene | 12580 |
| pRK5-FLAG-S6K1 | Addgene | 100509 |
| pRK7-HA-S6K1-F5A | Addgene | 8986 |
| pRK7-HA-S6K1-F5A-E389-R3A | Addgene | 8991 |
| pTRIP-CAG-mSVBP-mVASH2_X1-mCherry | CARSTEN JANKE | NA |
| pTRIP-CAG-hTTL-mscarlet | CARSTEN JANKE | NA |
| psPAX2 | Addgene | 12260 |
| pCMV-VSV-G | Addgene | 8454 |
| **Antibodies** |  |  |
| Acetylated α-tubulin | SigmaAldrich | T6793 |
| Detyrosinated α-tubulin | AbCam | ab48389 |
| GAPDH | Invitrogen | MA5-15738 |
| HA-Tag (6E2) | Cell Signalling Technology | 2367S |
| LAMP2 Antibody (H4B4) | Santa Cruz Biotechnology | sc-18822 |
| Mouse Anti Flag | Invitrogen | MA1-91878 |
| mTOR | Cell Signalling<br>Technology | 2983S |
| α-Tubulin | AbCam | ab18251 |
| P-P70 (S6K1) | Cell Signalling<br>Technology | 9205S |
| P70 (S6K1) | Cell Signalling<br>Technology | 2708S |
| P-4EBP1 | Cell Signalling<br>Technology | 2855S |
| 4EBP1 | Cell Signalling<br>Technology | 9644S |
| Occludin | Invitrogen | 33-1500 |
| Goat Anti Mouse AF 488 | Jackson Immuno<br>Research Laboratories Inc | 115-545-062 |
| Goat Anti Rabbit AF 488 Jackson (1mg) | Jackson Immuno<br>Research Laboratories Inc | 111-545-045 |
| Goat Anti Rat AF 488 Jackson (1mg) | Jackson Immuno<br>Research Laboratories Inc | 112-545-062 |
| Goat Anti Rat AF 647 Jackson (1mg) | Jackson Immuno<br>Research Laboratories Inc | 112-605-062 |
| Rabbit HRP-conj. secondary | Invitrogen | SA1-200 |
| Rat HRP-conj. secondary | Invitrogen | A18739 |
| Mouse HRP-conj. secondary | Invitrogen | SA1-100 |

### Immunofluorescence

Cell seeding was performed in 8-well Labtek chambered cover glass or confocal discs with a cell density of 15,000-40,000 cells per well with a seeding volume of 400 µl to 500 µl. Cells were fixed with fixation buffer (3% PFA & 1% GA, or 4% PFA, or absolute methanol at −20°C) for 10 min at room temperature or 5 min at −20°C or as mentioned otherwise. The autofluorescence of GA was quenched by using 1 mg/ml of NaBH₄ solution for 7 min at room temperature. Cells were blocked with 3% BSA containing 0.2% Triton X-100 or 0.2% saponin, or 4% BSA for 1-2 hr. Fixed cells were incubated in primary antibody for 1 hr at room temperature or overnight at 4°C, washed three times with washing buffer (0.2% BSA and 0.05% Triton X-100 or 0.2% Saponin) for 5 min each, incubated in secondary antibody for 40 min-1 hr, and washed three times with PBS for 5 min each. For double staining, either the primary and secondary antibodies were incubated together, or immunostaining for two different proteins was performed sequentially.

### Conventional microscopy

Fixed cell conventional imaging was done using a Nikon Ti2E microscope with a TIRF setup (100X oil immersion objective) using lasers of λ 488, 560, and 647 (MPB communications Inc.), and confocal imaging was done using a Zeiss LSM 700 confocal system with a 63X oil immersion objective using lasers of λ 488, 541, and 647.

### Colocalization analysis and quantification

For all the colocalization analysis (conventional and super-resolution) and quantification, including mTORC1 with lysosomes or mTORC1-lysosomes on tyrosinated/detyrosinated microtubules, the fluorescent signal from two different channels was manually estimated for overlap and scored.

### TFEB nuclear-to-cytoplasmic analysis

To quantify the percentage of cells with nuclear TFEB, EGFP TFEB confocal channel images from the mentioned conditions were manually analysed for higher nuclear intensity and scored as a cell positive for nuclear TFEB. To further quantitate accurately the abundance of nuclear TFEB, the raw integrated intensity of the unit area in the nuclear region is divided by that of the unit area in the cytoplasm to get the nuclear-to-cytoplasmic ratio.

### Lysotracker treatment

Cells were treated with Lysotracker Deep Red (Invitrogen, L12492) at 250 nM concentration for 20 min at RT. For the live tracking of lysosomes in starvation and restimulation, Lysotracker at 250 nM concentration was added to the starvation and restimulation media, respectively.

### Live imaging

For live imaging of lysosomes in BSC1 cells, cells were treated with lysotracker at respective conditions and were brought to the microscope at mentioned timepoints. Live imaging was performed in home-built TIRF microscope at a frame interval of 100 ms for 1 min.

For two-colour live imaging of lysosomes in Hap1 Raptor GFP cells, Hap1 cells, expressing stable endogenous level Raptor GFP, were treated with lysotracker, and live images were acquired with toggling of excitation lasers between 488 and 647 at a rate of 100 ms for 2 min.

For the two-colour live imaging for lysosome in Hap1 Raptor GFP cells throughout the restimulation time, the Leica Stellaris confocal microscope was used. Live movies were collected using an HC PL APO CS2 63x/1.40 oil objective, excited with a white light laser at 488 nm and 638 nm and detected using a Power HyD S detector. Laser scanning was done with a 1AU (95.6 µm) pinhole at a scan speed of 200 Hz.

### Zonal analysis of lysosomal distribution

The middle 10 out of the 600 frames of the lysosomal live movies were sub-stacked and z-projected to get the lysosomal position image. Zonal division of cellular cytosolic space was done using custom-written MATLAB code. The steps involve the generation of the cell boundary and nucleus using the respective DIC images, followed by the reduction of the cell boundary radially inward with respect to the nuclear centroid as its centre. The distance (in μm) that divides the cellular cytoplasmic region into three zones was used as the inward reduction distance. Thus, the cellular cytosolic space is divided into three distinct zones, perinuclear, middle, and peripheral, while preserving the cellular shape.

### Single particle tracking

Live movies of lysosomes were pre-processed with basic image corrections (in FIJI) to enhance the particle detection in single particle tracking, i.e., bleach correction with histogram matching and background subtraction with a rolling ball radius of 5 pixels. Then the movies were input to the tracking algorithm Trackit, a MATLAB-based tracking algorithm (Kuhn et al., 2021). The output trajectory was analysed with a custom-written Matlab analysis algorithm as mentioned in Mohan et al., (2019). Single lysosomal track displacement was analysed and displacement above the threshold value of 1 µm is considered to be mobile lysosome tracks, while displacement below 1 µm is considered to be non-mobile lysosomes. The threshold value of 1 µm was chosen because of two reasons. 1) The mean displacement of lysosome in control untreated conditions was found to be 1 µm. 2) The average diameter of lysosomes varies from 0.5 to 0.8 µm.

For the track mean speed analysis of lysosome with and without lysosomes, live movies of the Raptor GFP and Lysotracker channel were obtained from TIRF microscopy and pre-processed as mentioned in the above section. Then the lysosomes were manually annotated for the presence or absence of Raptor GFP and tracked in TrackMate, an imageJ tracking plugin (Ershov et al., 2022). The mean speed of the lysosome with and without mTORC1 were compared.

For live movies of lysosome in Hap1 Raptor GFP cells throughout the restimulation time, they were pre-processed as mentioned in the previous section and/or image smoothened for better detection of Raptor GFP intensity on the lysosome (Lysotracker). Then the movies were carefully observed in order to find the event of increasing mTORC1 (Raptor GFP) intensity on lysosome over restimulation time. Those lysosomes were manually annotated and tracked over time in TrackMate.

### Anterograde and retrograde transport classification

Classification and quantification of transport directionality was done using a custom-written Python-based algorithm. The start and end coordinates of the lysosomal trajectories were connected to form displacement vectors representing the displacement. The cell boundary and the nuclear boundary were fed to the algorithm. The non-mobile lysosomal tracks were filtered out. Two directional vectors were formed, from the start point of the track to the point on the cellular boundary (directional vector 1) and nuclear boundary (directional vector 2) that were nearest to the end point. The difference between the dot product of the displacement vector with directional vector 1 and the dot product with directional vector 2 was calculated. If this difference was greater than 0.68, the track would be classified as retrograde (towards the nucleus). If the difference was less than - 0.68, the track would be classified as anterograde (towards cell boundary), and the rest are classified as indecisive.

### Microtubule polymerization quantification

Singular, clearly moving microtubule EB3 were cropped from raw time-lapse image sequences in ImageJ to minimize background crosstalk and noise. Kymographs were generated using the KymographClear plugin where, for each microtubule end (EB3), a segmented line ROI was drawn along the microtubule growth path, and the plugin’s “Kymograph Generation” function was applied to produce kymographs out of which Kymograph_filtered_forward.tif was utilized for further analysis. Microtubule growth speed was quantified by tracing a segmented line along the kymograph trajectory and computing the slope (Δdistance/Δtime) of each linear segment. Segment-wise slopes were then averaged to obtain the final growth speed for that microtubule.

### Super-resolution Structured illumination microscopy

SIM imaging was performed in ZEISS Elyra 7 (Lattice SIM Carl Zeiss Ltd.). Images were acquired by exciting with λ 488, 561, and 642 lasers and collected using a Plan-Apochromat 63x/1.4 oil immersion objective with an in-built filter cube set for the respective fluorescent channel with optical sectioning (Z stack) in 3D. SIM image processing was performed with the manufacturer’s in-built software in 3D standard mode. SIM imaging was performed in a Zeiss Elyra microscope at Sathi, IIT Delhi Sonipat Campus.

### SIM image processing and MT percent quantification

SIM images (two MT channels) were maximum or sum intensity projected along the z-axis (Z-project) and converted to 16-bit images. Then, to ensure that the level of detyrosinated MT is quantified over the total MT population, both tyrosinated and detyrosinated MT images were combined to get the total MT population. Subsequently, total MT and detyrosinated MT images were auto-thresholded using the Isodata or moments algorithm in Fiji. Then, to get the fraction of detyrosinated MT population, the overlap fraction (Mander’s coefficient) of detyrosinated MT over the total MT image was calculated using the JACop plugin in Fiji (https://imagej.net/plugins/jacop) and multiplied by 100 to get the percentage of detyrosinated MT population. The same method was followed to quantify acetylated MT population. Only for the conditions where the detyrosinated MT are very minimal for thresholding and quantification, the percentage of tyrosinated MT is calculated and subtracted from 100 to get the percentage of detyrosinated MT.

### STORM Imaging

STORM imaging was performed using a home-built STORM microscope setup. A 647 nm or a 488 nm laser (MPB Communications) was used to excite the Alexa Fluor 647 (Invitrogen) or Alexa Fluor 488, respectively, and a 405 nm laser (Obis; Coherent) was used for activation. Emitted light was collected with a Nikon 100x oil immersion TIRF-SR objective (NA 1.49) and laser quad band set with emission filter (TRF89902-EM ET - 405/488/561/647 nm Laser Quad Band Set for TIRF applications; Chroma). Images were captured using an EM-CCD camera at an exposure time of 20 milliseconds per frame. A total of 60000 frames were acquired to reconstruct a single STORM image. Two-colour STORM was performed sequentially. First STORM images were acquired for the first protein of interest stained with Alexa Fluor 647, then the fluorescence signals were quenched by treating the samples with 1 mg/mL of sodium borohydride (sodium borohydride, Sigma 213462) for 7 min at room temperature, followed by staining for the second protein of interest with AF 647 and performing STORM imaging of the second protein. STORM images were reconstructed and rendered using custom-written software (Insight3, provided by B. Huang, University of California, San Francisco, CA) as previously described in Verdeny-Vilanova et al., (2017). Multi-channel STORM images were corrected for chromatic shifts and mechanical drifts and were spatially aligned using localisations of fiduciary markers (TetraSpeck microspheres, Invitrogen T7279) across the two different fluorescence channels as described previously in Bálint et al., (2013).

For STORM imaging, the stained cells were incubated during acquisition in an imaging buffer containing 50mM cysteamine, 3% [Vol/Vol] Oxyfluor (Sigma-SAE0059), and 20% [Vol/Vol] Sodium DL-lactate (Sigma-L1375) in PBS at pH 8.

### Correlative Live STORM imaging (CLSM)

Live imaging of mTOR YFP in the BSC1 mTOR YFP cell line was performed in a home-built TIRF microscope at a frame interval of 100 ms for 2 min and immediately fixed with PFA GA on top of the microscope. Then, immunostaining for tyrosinated or detyrosinated microtubules was performed on this sample kept on the microscope stage carefully without disturbing the position. Thereafter, STORM imaging for tyrosinated or detyrosinated microtubules was carried out as mentioned in the previous section. STORM images were reconstructed, corrected for chromatic shifts and mechanical drifts, and spatially aligned to the live movie using localisations of fiduciary markers (TetraSpeck microspheres, Invitrogen T7279) as described previously in Bálint et al., (2013).

### Immunoblotting

Cells were lysed with RIPA buffer supplemented with protease inhibitor cocktail (Promega) and phosphatase inhibitor cocktail (GbioSciences-786-870), incubated on ice for 30 min, and vortexed 4 times at 15 min intervals. Then the samples were sonicated and centrifuged at 14,000 RPM for 20 min at 4°C. The supernatant was collected and estimated for protein concentration. Cell lysates were reduced in 2-mercaptoethanol-containing Laemmli buffer and loaded at equal volume in 10% and 12% polyacrylamide gels of 1-mm thickness. After the SDS-PAGE run, the proteins from the gel are transferred to the polyvinyl difluoride membrane (Millipore; Immobilon-FL, 0.45-µm thickness) using wet transfer in Towbin buffer. Transfer voltage was maintained at 100V for 90 min under ambient condition with an ice block inside and ice surrounding the tank. After the transfer, the membrane was stained with Ponceau to check the transfer and equal loading. Then the membranes were cut for necessary protein bands and blocked for 1-2 hr at room temperature in 3% BSA or 5% NFDM with 0.1% Tween 20. The primary antibody was incubated overnight in 3% BSA with 0.1% Tween 20. The membranes were washed with TBST for 10 min for 2 times and were incubated in the secondary antibody conjugated with HRP for 1 hr at room temperature protected from light. The membranes were washed in TBST for 10 minutes, three times. The proteins in the membrane were detected using chemiluminescent HRP substrate (SuperSignal™ West Pico PLUS Chemiluminescent Substrate, Invitrogen-34580, and Clarity Max Western ECL Substrate, 100 ml Biorad-1705062) in the Biorad gel documentation system (Gel Doc XR+). The images are acquired with the manufacturer-provided imaging software Image Lab.

### Image Densitometry

The acquired blot images were quantified in the same Image Lab software. The protein band intensity value was background subtracted and normalised to the background subtracted intensity value of the loading control (GAPDH) for each blot.

### Wound healing assay

22RV1 WT and VASH2 mCherry cells were seeded in a 12-well plate at a seeding density of cell/well. After 48 hr of 70-90% confluency, cells were washed gently and replenished with fresh media and scratch were made with sterile 200 µL pipette tip. Then the image of the 0^th^ hr scratch was acquired using an Olympus phase contrast microscope with a 10x objective, and the coordinates were noted for further imaging. Then for a 120 hr time period, scratch site images were acquired every 12 hr. The wound closure area and migration rate were estimated using the ImageJ plugin, Wound healing size tool (Suarez-Arnedo et al., 2020).

### Statistical analysis

All data are represented in mean ± SD, and the number of replicates is mentioned in the figure legends. For all the experiments with two groups, the unpaired student’s t-test was used. For all the experiments with more than two groups and all Western blot fold change quantification, one-way ANOVA with Tukey test was applied. For wound area closure with respect to time, Two-way mixed RM-ANOVA (Time within, Group between) with GG correction when Mauchly’s test indicated sphericity violation; α=0.05. Simple effects reported with Tukey-adjusted pairwise tests. Significance in difference were marked as *P<0.05, **P<0.005, ****P<0.00005.

## Supporting information

Figure 1C- Lysosomal movements at all nutrient condition time points

Figure1G- Lysosome with mTORC1 movie

Figure1G- Lysosome without mTORC1 movie

Figure 2H- Lysosomal movement under various nutrient conditions

Figutre 3C- mTORC1 movement on Tyrosinated MT

Figutre 3C- mTORC1 movement on Detyrosinated MT

Figure 5A- Lysosomal movement on Tyro/Detyro MT level modulation

Figure S1C- mTORC1-lysosomal recruitment and movement on AA FBS Restimulation

Figure S2I- Microtubule growth on Various nutrient conditions

## Data availability statement

The data set studied and investigated, and algorithm used during the study are accessible from the corresponding author upon reasonable request.

## Acknowledgments

We thank Prof. Melike Lakadamyali (University of Pennsylvania, Philadelphia, PA) for supporting the research with reagents and software. We thank Prof. Nicholas T Ktistakis (Babraham Institute, Cambridge) and Prof. Bushra Ateeq (IIT Kanpur, India) for generously providing cell lines. We thank Prof. Carsten Janke (Institut Curie, France), Prof. Minhaj Sirajuddin (InStem, Bengaluru, India), and Prof. Sai Prasad Pydi (IIT Kanpur, India) for providing plasmids. We thank Prof. Suresh Kumar (IIT Kanpur, India) for providing reagents, facility and guidance for stable clone generation. We Thank Prof. Pradip Sinha (IIT Kanpur, India) for providing antibody. We thank Prof. Saravanan Matheshwaran (IIT Kanpur, India) for providing facilities and reagents for microbial work.

We thank the Sophisticated Analytical & Technical Help Institutes (SATHI, IITD; DST project Sanction Number: RD/SATHI/NEW-CENTERS/IIT-DELHI/2019 (G)) for providing access to the Structure Illumination Microscopy (SIM) and the Central Research Facility – IIT Delhi. We also extend our thanks to Mr. Taseen Ahmad for assisting in SIM imaging.

J.A. expresses sincere gratitude towards the MHRD and the Indian Institute of Technology, Kanpur, India for financial support. J.A. gratefully acknowledges Shefali T, Deepak M. Khushalani, and Sanket Patil in helping establish the laboratory at IIT Kanpur, India. J.A. acknowledges Jemina J for generation of AG/RG analysis algorithm used in the study. J.A. also acknowledges Ms. Deepa Ajnar for the support in WB and Stable clone generation, Ms. Sushmita Kundu and Mr. Umar Khalid Khan for helping with cancer cell lines and inputs. J.A. expresses sincere acknowledgement towards Dr. Vijesh K R for support in analysis, Dr. Rachita Battacharya, Mr. Karthikeyan, Mr. Hariharan VC for helping with reagents and the technical support.

## Declaration of interest statement

The author declares that there is no conflict of interest or any personal relationship that could have influenced the work reported in this paper.

## Notes

### Competing Interest Statement

The authors have declared no competing interest.

