## Supplementary Figures for "mTORC1 signaling modulate microtubule tyrosination/detyrosination status to regulate lysosome dynamics"

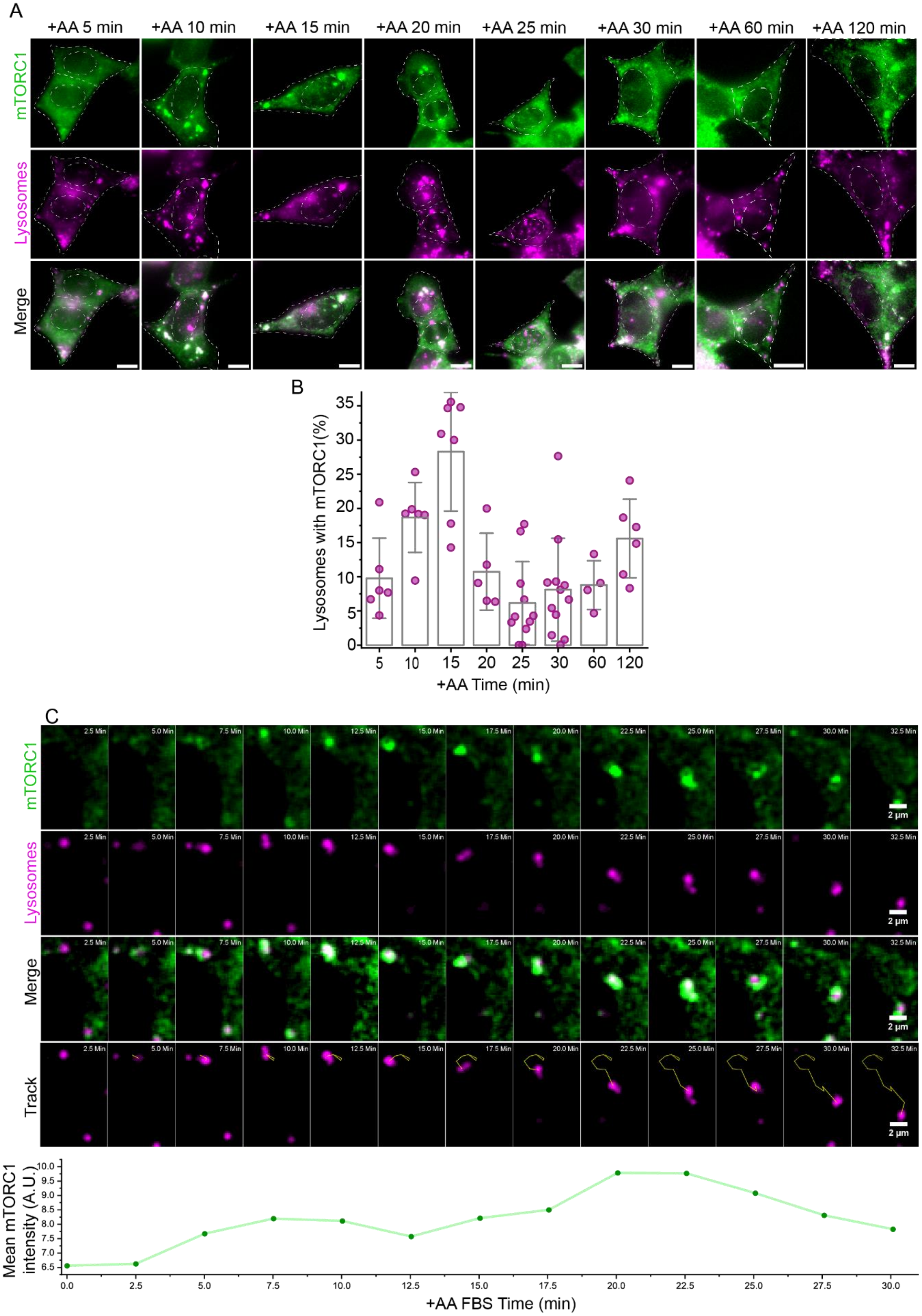

**Figure S1 (A)** Two-colour TIRF microscopy of endogenous mTOR (green) and LAMP2 (lysosomes, magenta) in Hap1 Raptor GFP cells starved for 1 hours or starved and restimulated with amino acids for indicated time points. **(B)** Quantification of lysosomes positive for mTOR signal. Starved cells show minimal mTOR-lysosome colocalization, which increases upon nutrient restimulation, peaking at 15 minutes, followed by a decline at 20 minutes. **(C)** Live-cell confocal imaging of HAP-1 cells expressing endogenous RAPTOR-GFP (mTORC1) labelled with Lysotracker. Cells were starved and restimulated as in **(A)** and imaged for Raptor GFP and Lysosome at every 2.5 min of restimulation time till 32.5 min of restimulation. Lysosome gains higher motility as the localization of mTORC1 increases at $15^{th}$ to $17^{th}$ min of restimulation. The bottom plot shows the mean grey value of the RAPTOR GFP signal in the ROI shown, representing the increase in the lysosomal localization of mTORC1 with restimulation time. The scale bar for A is 10 μm and C is 2 μm.

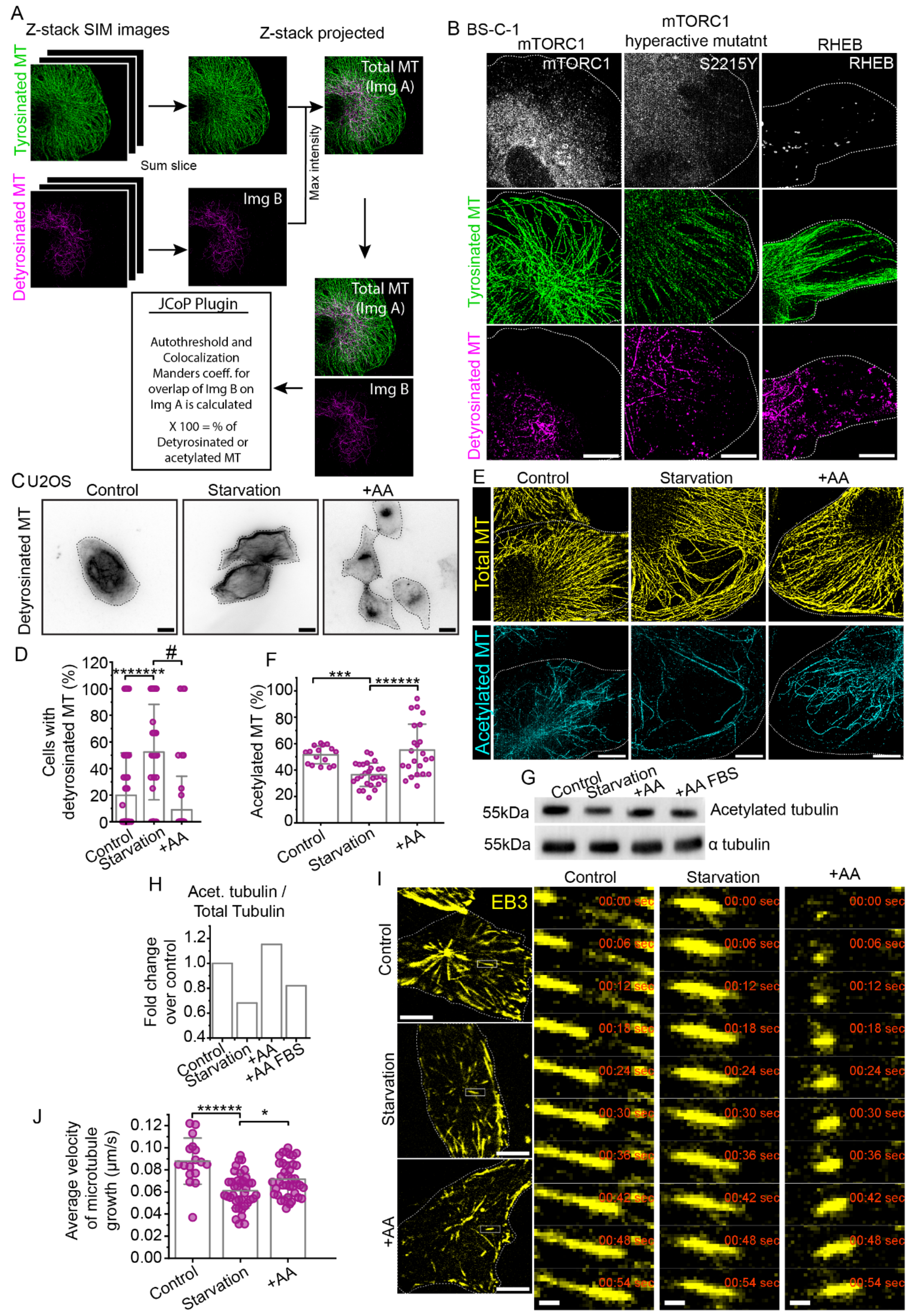

A
Z-stack SIM images
Z-stack projected
Tyrosinated MT
Detyrosinated MT
Sum slice
Max intensity
Total MT (Img A)
Img B
JCoP Plugin
Autothreshold and Colocalization Manders coeff. for overlap of Img B on Img A is calculated
X 100 = % of Detyrosinated or acetylated MT
B BS-C-1
mTORC1
mTORC1 hyperactive mutatnt
RHEB
mTORC1
S2215Y
RHEB
C U2OS
Control
Starvation
+AA
Detyrosinated MT
D
Cells with detyrosinated MT (%)
#
*******
E
Total MT
Acetylated MT
F
Acetylated MT (%)
***
******
G
Control
Starvation
+AA
+AA FBS
55kDa
Acetylated tubulin
α tubulin
H
Acet. tubulin / Total Tubulin
Fold change over control
I
EB3
00:00 sec
00:06 sec
00:12 sec
00:18 sec
00:24 sec
00:30 sec
00:36 sec
00:42 sec
00:48 sec
00:54 sec
J
Average velocity of microtubule growth (µm/s)
******
*

**Figure S2** **(A)** Schematics of SIM image quantification pipeline **(B)** Two colour SIM microscopy of tyrosinated (green) and detyrosinated (magenta) microtubule in BS-C-1 cells transiently over expressed with mTORC1 WT or mTORC1 hyperactive mutant (S2215Y) or RHEB **(C)** TIRF microscopy of detyrosinated microtubules (grey) in U2OS cells untreated or starved for 2 hr or starved and restimulated with 10X AA for 30 min **(D)** Manual scoring of percentage of cells with detyrosinated microtubule in the mentioned conditions in **(C)**, shows the increase in the population of cells with detyrosinated microtubules under starvation condition **(E)** Two colour SIM microscopy of acetylated (cyan) and alpha (yellow) microtubule in BS-C-1 cells untreated or starved for AA and FBS for 2 hr or starved and restimulated with AA for 15 min **(F)** Quantification of acetylated microtubules shows decrease in the percentage of acetylated microtubule in starvation and increase under AA restimulation **(G)** Immunoblotting for acetylated and alpha tubulin in BS-C-1 cells starved for 2 hr or starved and restimulated with AA or AA and FBS for 20 min **(H)** Densitometric quantifications of protein bands under different conditions mentioned in **(G)** show the decrease in the acetylated tubulin on starvation. **(I)** Live-cell TIRF imaging of End binding protein (EB3) in BS-C-1 cells untreated or starved for 2 hr or starved and restimulated with AA for 20 min.**(J)** Average velocity of microtubule quantification by kymograph analysis of microtubule growth showing the reduced growth dynamics of microtubule in starvation condition. The bar in all plots represents the mean, error bars represent SD; $^*P<0.05$, $^{**}P<0.005$, $^{****}P<0.00005$. The scale bar for all the images is 10 μm.

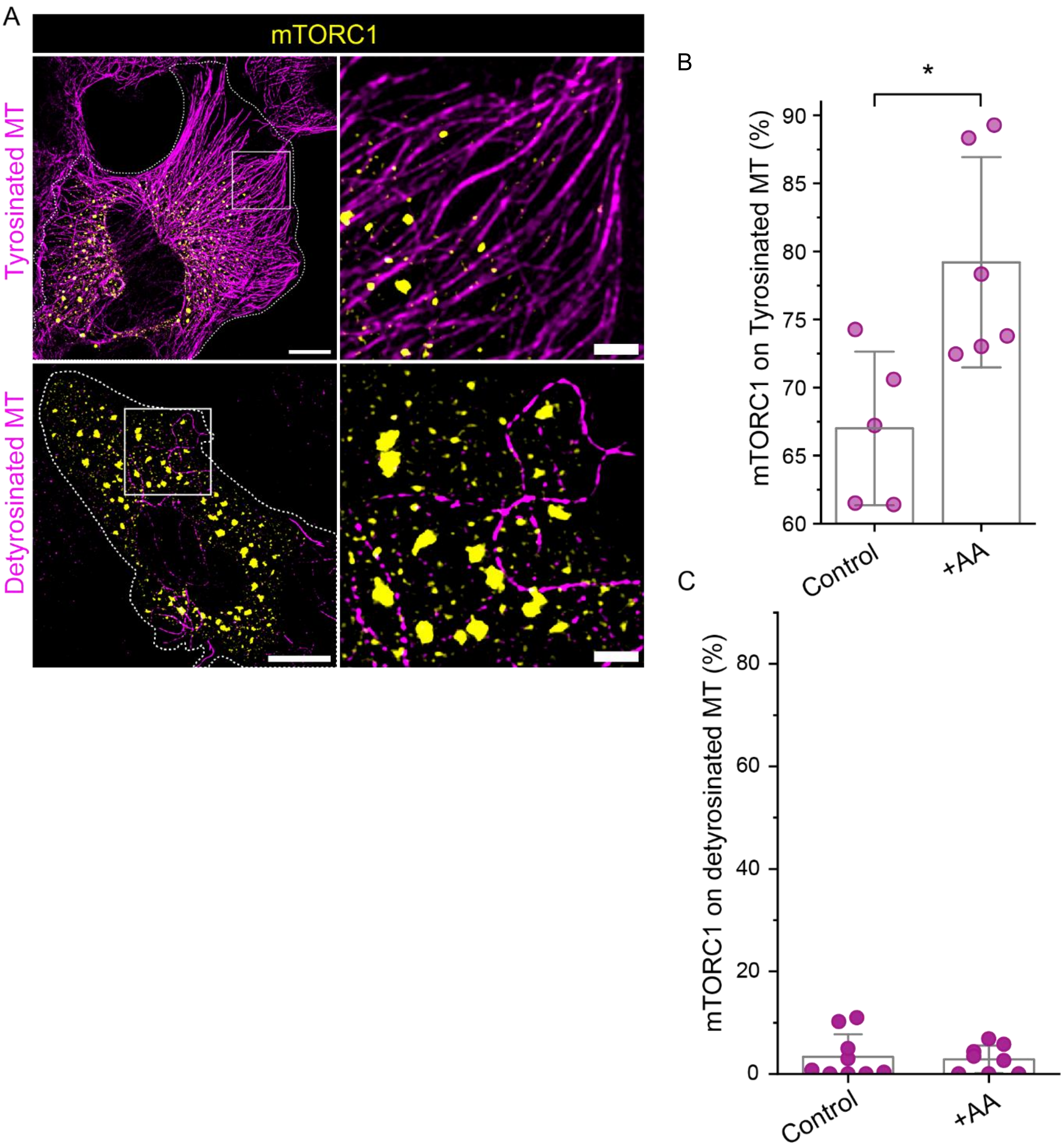


**Figure S3. (A)** Two-colour SIM microscopy of tyrosinated or detyrosinated microtubule (magenta) and over expressed mTORC1 (yellow) in BS-C-1 cells. Manual scoring of mTORC1-positive lysosomes on **(B)** tyrosinated or **(C)** detyrosinated microtubules shows an increase in the percentage of mTORC1-positive lysosomes localising to tyrosinated microtubules on AA FBS restimulation and minimal localisation of mTORC1-positive lysosomes on detyrosinated microtubules irrespective of nutrient condition. The bar in the plot represents the mean, and error bars represent SD; *P<0.05, **P<0.005, ****P<0.00005. The scale bar for **(A)** is 10 μm and 2 μm for inset zoom images

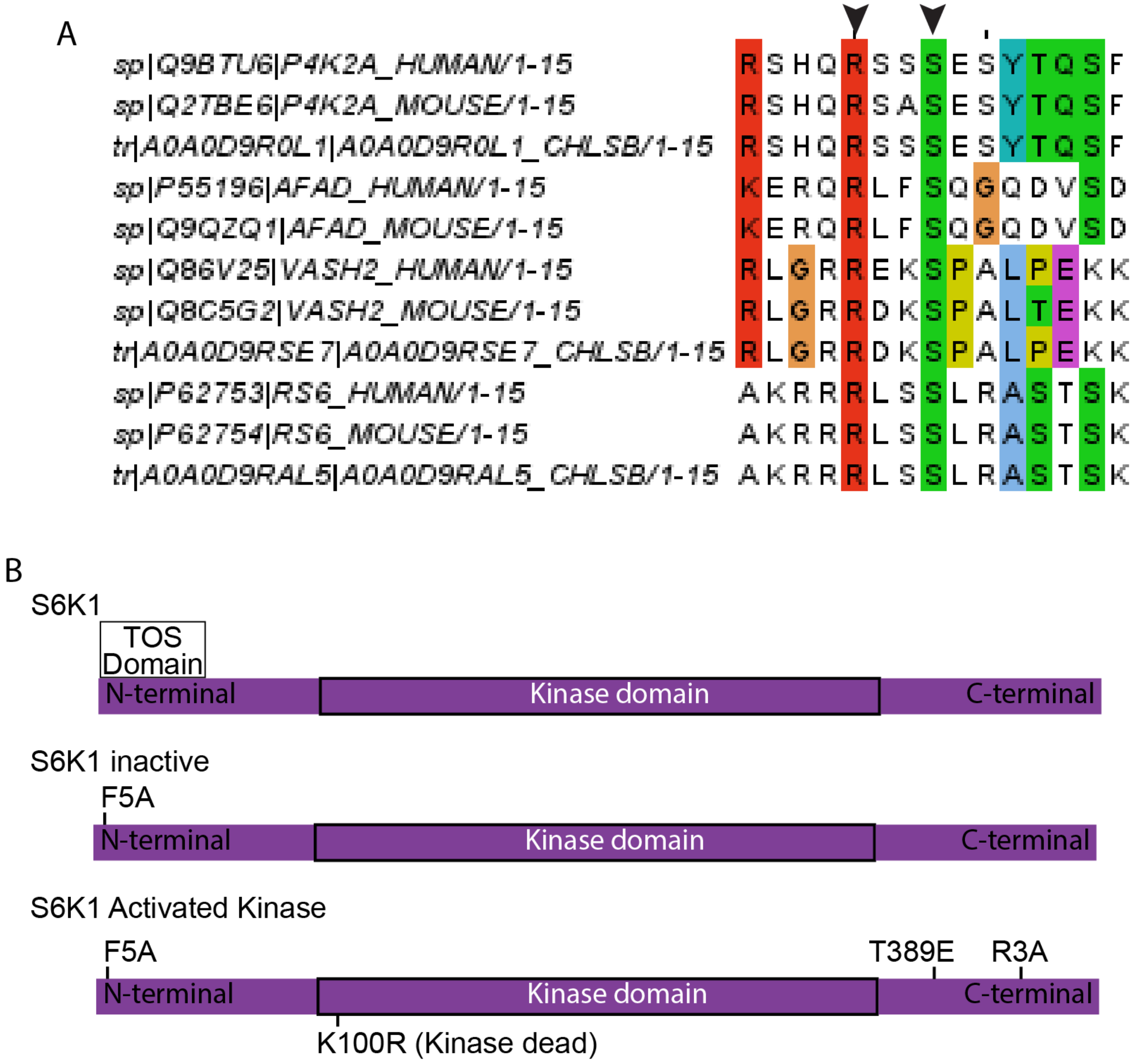


**Figure S4. (A)** Multiple sequence alignment of the putative S6K1 phosphorylation motif (RXXS) in VASH2 with the reported site in known S6K1 substrates across mammalian species. **(B)** Schematic outline of S6K1 variants used to modulate activity genetically.

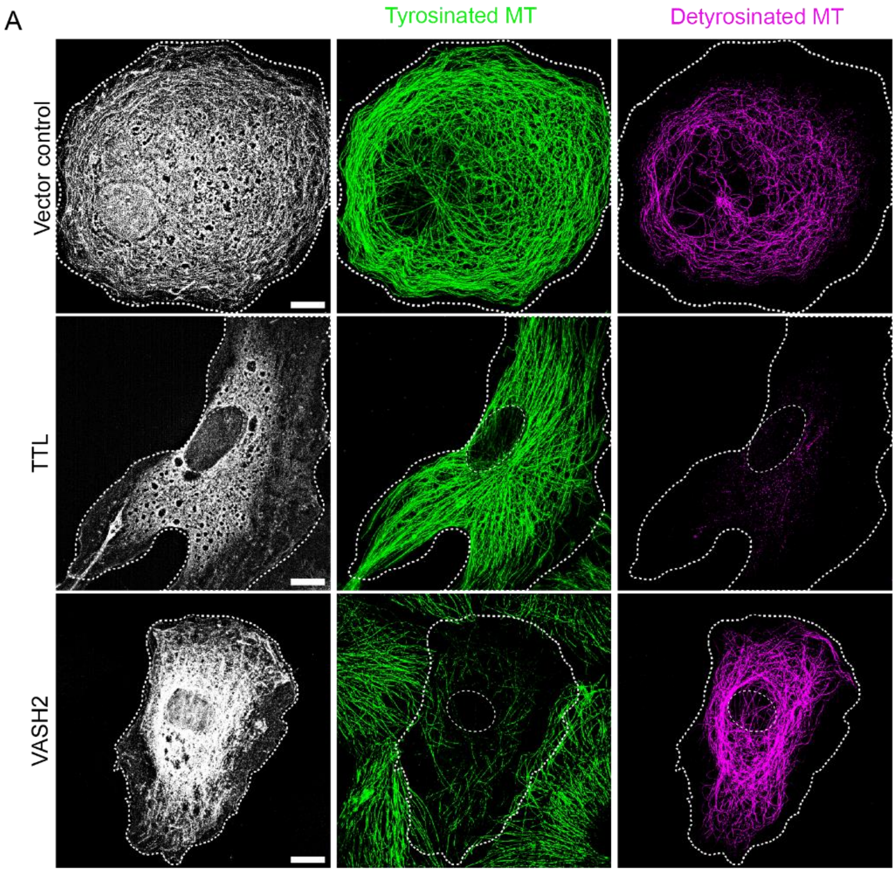


**Figure S5. (A)** SIM microscopy of tyrosinated (green) and detyrosinated (magenta) microtubules in BS-C-1 cells transfected with vector control or TTL mscarlet or VASH2 mCherry reveal the increase of tyrosinated microtubules and detyrosinated microtubules on TTL and VASH2 overexpression, respectively.

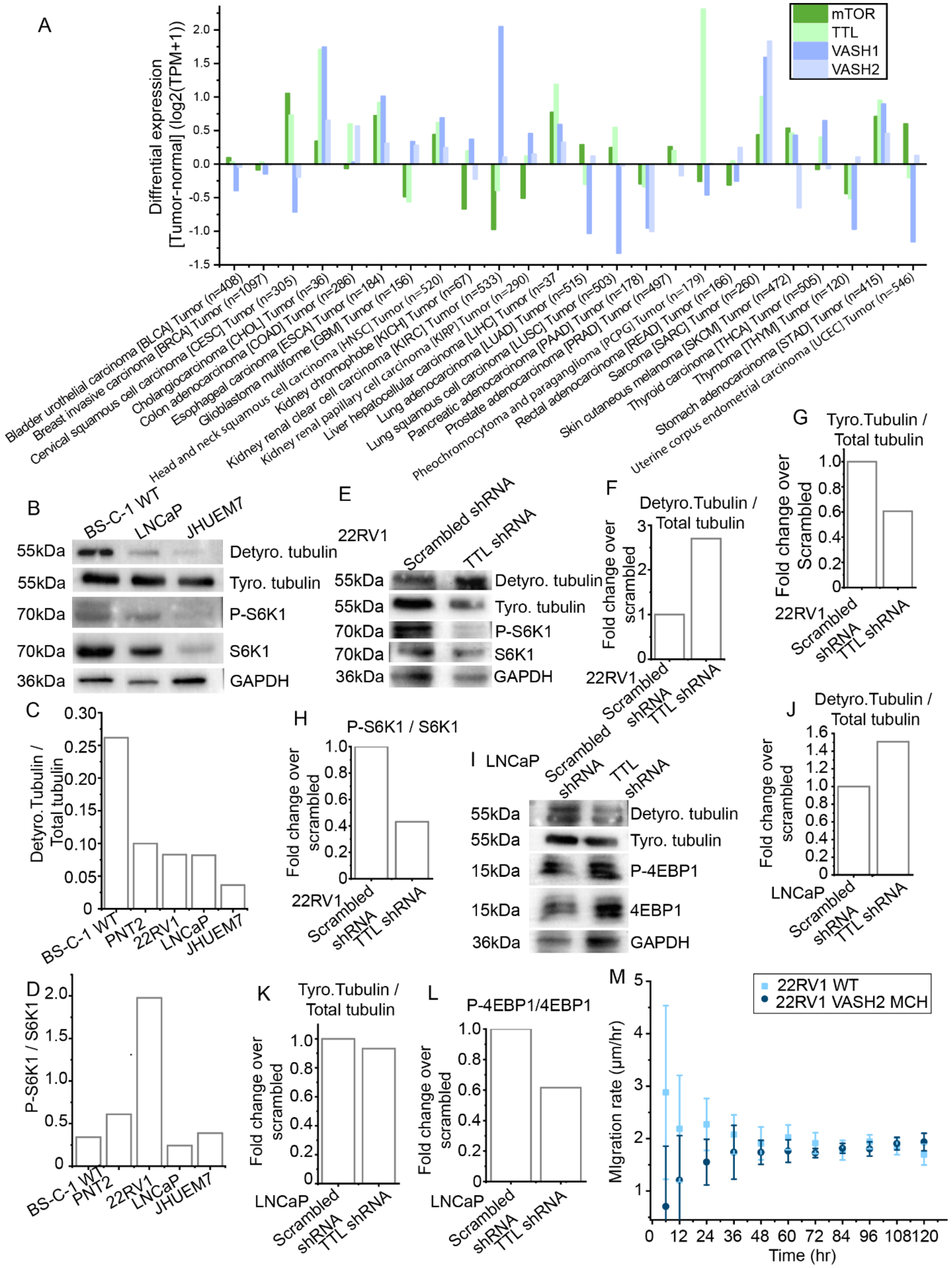
A
Diffrential expression [Tumor-normal] (log2(TPM+1))
mTOR
TTL
VASH1
VASH2
Bladder urothelial carcinoma [BLCA] Tumor (n=408)
Breast invasive carcinoma [BRCA] Tumor (n=1097)
Cervical squamous cell carcinoma [CESC] Tumor (n=305)
Cholangiocarcinoma [CHOL] Tumor (n=36)
Colon adenocarcinoma [COAD] Tumor (n=286)
Esophageal carcinoma [ESCA] Tumor (n=184)
Glioblastoma multiforme [GBM] Tumor (n=156)
Head and neck squamous cell carcinoma [HNSC] Tumor (n=520)
Kidney chromophobe [KICH] Tumor (n=67)
Kidney renal clear cell carcinoma [KIRC] Tumor (n=533)
Kidney renal papillary cell carcinoma [KIRP] Tumor (n=290)
Liver hepatocellular carcinoma [LIHC] Tumor (n=37
Lung adenocarcinoma [LUAD] Tumor (n=515)
Lung squamous cell carcinoma [LUSC] Tumor (n=503)
Pancreatic adenocarcinoma [PAAD] Tumor (n=178)
Prostate adenocarcinoma [PRAD] Tumor (n=497)
Pheochromocytoma and paraganglioma [PCPG] Tumor (n=179)
Rectal adenocarcinoma [READ] Tumor (n=166)
Sarcoma [SARC] Tumor (n=260)
Skin cutaneous melanoma [SKCM] Tumor (n=472)
Thyroid carcinoma [THCA] Tumor (n=505)
Thymoma [THYM] Tumor (n=120)
Stomach adenocarcinoma [STAD] Tumor (n=415)
Uterine corpus endometrial carcinoma [UCEC] Tumor (n=546)
B
BS-C-1 WT
LNCaP
JHUEM7
55kDa Detyro. tubulin
55kDa Tyro. tubulin
70kDa P-S6K1
70kDa S6K1
36kDa GAPDH
C
Detyro.Tubulin / Total tubulin
BS-C-1 WT
PNT2
22RV1
LNCaP
JHUEM7
D
P-S6K1 / S6K1
BS-C-1 WT
PNT2
22RV1
LNCaP
JHUEM7
E
22RV1
Scrambled shRNA
TTL shRNA
55kDa Detyro. tubulin
55kDa Tyro. tubulin
70kDa P-S6K1
70kDa S6K1
36kDa GAPDH
F
Detyro.Tubulin / Total tubulin
Fold change over scrambled
22RV1
Scrambled shRNA
TTL shRNA
G
Tyro.Tubulin / Total tubulin
Fold change over Scrambled
22RV1
Scrambled shRNA
TTL shRNA
H
P-S6K1 / S6K1
Fold change over scrambled
22RV1
Scrambled shRNA
TTL shRNA
I
LNCaP
Scrambled shRNA
TTL shRNA
55kDa Detyro. tubulin
55kDa Tyro. tubulin
15kDa P-4EBP1
15kDa 4EBP1
36kDa GAPDH
J
Detyro.Tubulin / Total tubulin
Fold change over scrambled
LNCaP
Scrambled shRNA
TTL shRNA
K
Tyro.Tubulin / Total tubulin
Fold change over scrambled
LNCaP
Scrambled shRNA
TTL shRNA
L
P-4EBP1/4EBP1
Fold change over scrambled
LNCaP
Scrambled shRNA
TTL shRNA
M
22RV1 WT
22RV1 VASH2 MCH
MIgration rate (μm/hr)
Time (hr)

**Figure S6. (A)** Differential expression analysis of mTOR, TTL, VASH1/2, HDAC6 in tumour sample with respect to normal sample collected from UALCAN database showing the expression correlation between mTOR with TTL or VASH1/2 in multiple cancer types. **(B)** Immunoblotting for tyrosinated and detyrosinated tubulin along with mTORC1 substrate S6K1 (phosphorylated and total) in BS-C-1, LNCaP, and JHUEM7 **(C-D)** Densitometric quantifications of protein bands in **(B and Figure 6B)** show the increase in detyrosinated tubulin and decrease in mTORC1 activity represented by the phosphorylated level of S6K1. **(E and I)** Immunoblotting for tyrosinated and detyrosinated tubulin along with mTORC1 substrates S6K1 or 4EBP1 (phosphorylated and total) in 22RV1 and LNCaP cells under TTL knockdown **(F-H, J-L)** Densitometric quantifications of protein bands in **(E and I)** show the decrease in tyrosinated tubulin and mTORC1 activity represented by the phosphorylated level of S6K1 or 4EBP1 **(M)** Quantification of wound closure area for **Figure 6G** showing decreased migration rate in VASH2 mCherry overexpressed 22RV1 cells.
